# Autocrine CXCL1-CXCR2 Signaling Mediates Leptomeningeal Resistance to Radiation Therapy

**DOI:** 10.1101/2025.02.20.639389

**Authors:** Ahmed M. Osman, Jan Remsik, Jenna Snyder, N. Ari Wijetunga, Branavan Manoranjan, Ana Rita Nobre, Morgan Freret, David Guber, Kiana Chabot, Sofia Piedrafita-Ortiz, Xinran Tong, Helen Wang, Min Jun Li, Andrew J. Dunbar, Ross L. Levine, Jonathan T. Yang, Adrienne Boire

## Abstract

Leptomeningeal metastasis (LM) is a fatal neurological complication of cancer. Proton craniospinal irradiation (pCSI) has emerged as a promising life-prolonging intervention for LM patients, but the response to this treatment varies. Here, we aimed to characterize the molecular basis of pCSI resistance and response. Proteomic analysis of CSF collected from LM patients at baseline (before pCSI), and at multiple time points post-treatment, identified the CXC-motif chemokine, CXCL1, as associated with LM growth. Higher CXCL1 levels in the CSF prior to pCSI correlated with worse response to this treatment. To define the role of CXCL1 in LM, we established syngeneic mouse models of LM-CSI. We found that both metastatic cancer and host cells generate CXCL1. Genetic interruption of *Cxcl1* expression in metastatic cancer, but not host cells, impaired cancer cell growth within the leptomeninges. Moreover, a subset of LM cancer cells expressed Cxcr2, the primary receptor for Cxcl1, and this population was enriched over time in the leptomeninges. Transcriptomic profiling of this rare population revealed an enrichment in pathways implicated in cell cycle progression. Finally, interruption of Cxcl1-Cxcr2 signaling with intrathecally-delivered Cxcr2 antagonist hampered LM growth and sensitized the cells to CSI. Our results demonstrate that the Cxcl1-Cxcr2 signaling axis mediates LM growth, and identifies a potential actionable intervention to improve response to pCSI and halt LM progression.

**One Sentence Summary:** CXCL1-CXCR2 axis is a potential actionable therapeutic target to halt leptomeningeal metastasis progression and enhance response to craniospinal irradiation.

## Introduction

Leptomeningeal metastasis (LM), also known as leptomeningeal disease, is a fatal neurological complication of cancer characterized by growth of metastatic cancer cells within the cerebrospinal fluid (CSF)-filled leptomeningeal compartment encasing the brain and the spinal cord. LM occurs in 5-20% of patients with solid tumors, and is most likely to occur secondary to breast cancer, lung cancer, and melanoma (*1, 2*). Cancer cells enter the leptomeningeal compartment through a variety of means, including the choroid plexus (*3–5*). Upon entrance into the leptomeningeal compartment, cancer cells provoke an intense inflammatory response characterized by a host of cytokines (*6*). We have previously found that metastatic cancer cells exploit this inflammation to perturb the blood-CSF-barrier, and produce a high-affinity iron collection and uptake system (LCN2 and SLC22A17) to support their growth (*3, 7*). What other mechanistic purposes inflammatory signaling might serve in the leptomeningeal space and LM metastatic capacity remain under-explored (*8*).

Clinically, cancer cells within the leptomeningeal compartment adopt two phenotypic states: plaques of tumor adherent to the brain and spinal cord, best appreciated by magnetic resonance imaging (MRI); and the free-floating cells within the spinal fluid, detected via CSF cytology (*1, 2, 9*). After diagnosis, untreated LM patients succumb within 4 - 6 weeks after diagnosis, or 2 - 5 months when treated with the current available therapies (*1, 2, 10*). Recently, proton craniospinal irradiation (pCSI) has been demonstrated as a life-prolonging intervention in recent clinical trials (NCT03520504 and NCT04343573) (*11, 12*). While a cohort of pCSI-treated patients survived over 18 months (responders), others did not and succumbed to their illness within 3 months post-treatment (non-responders). The molecular basis of pCSI response and resistance in LM remains unknown. In this study, we performed proteomic analyses on serially collected CSF from LM patients at baseline (before pCSI) and after treatment. We then developed preclinical immune-competent LM-CSI models to define signaling programs implicated in pCSI resistance and response that can be exploited to maximize pCSI benefit.

## Results

### CSF CXCL1 levels are elevated in LM and portend poor response to pCSI

While pCSI was demonstrably superior to photon involved-field radiotherapy for control of LM (*11*), a cohort of patients did not respond to the treatment and demonstrated LM progression despite the treatment (Fig. 1A). The LM microenvironment is nutrient-sparse (*13*), and the presence of cancer cells within this space triggers a florid inflammatory response (*6, 7, 14*). Despite this particularly harsh microenvironment, cancer cells persist, grow, and resist therapies. We hypothesized that the byproduct of the signaling events governing LM growth could be reflected in the CSF. Accordingly, we assayed the cell-free fractions of serially collected CSF from patients enrolled in our clinical trial (NCT03520504), obtained before treatment (baseline), and at 1-, 3-, and 6-months post-treatment, by targeted proteomic analysis using proximity ligation assay (fig. S1A). Given the enriched immune response in the CSF of LM-harboring patients, and the crucial role inflammatory signals play to mediate LM growth (*6, 7*), we focused on 92 proteins related to inflammation. We found that pCSI resulted in a significant reduction in the levels of the chemokine CXCL1 one month post-treatment compared to baseline, and these levels were maintained (Fig. 1B). These results were confirmed by conventional ELISA (fig. S1B). We next asked if the reduced levels of CXCL1 post-treatment were due to the effect of irradiation regardless of LM, or if this might be secondary to the presence of LM alone. To investigate this, we extended our patient cohort and analyzed the levels of CXCL1 in the CSF of non-irradiated patients with a variety of solid primary tumors, both with and without LM. We found that LM led to a significant increase in CXCL1 (Fig. 1C, and fig. S1C). Taken together, these results demonstrate that the presence of LM leads to elevated CXCL1 levels in the CSF, and pCSI reduces CSF CXCL1 levels. In patients treated with pCSI, we found that higher CXCL1 levels before treatment were associated with worse CNS progression-free survival, but did not significantly alter overall survival (Fig. 1, D and E).

**Figure 1:**
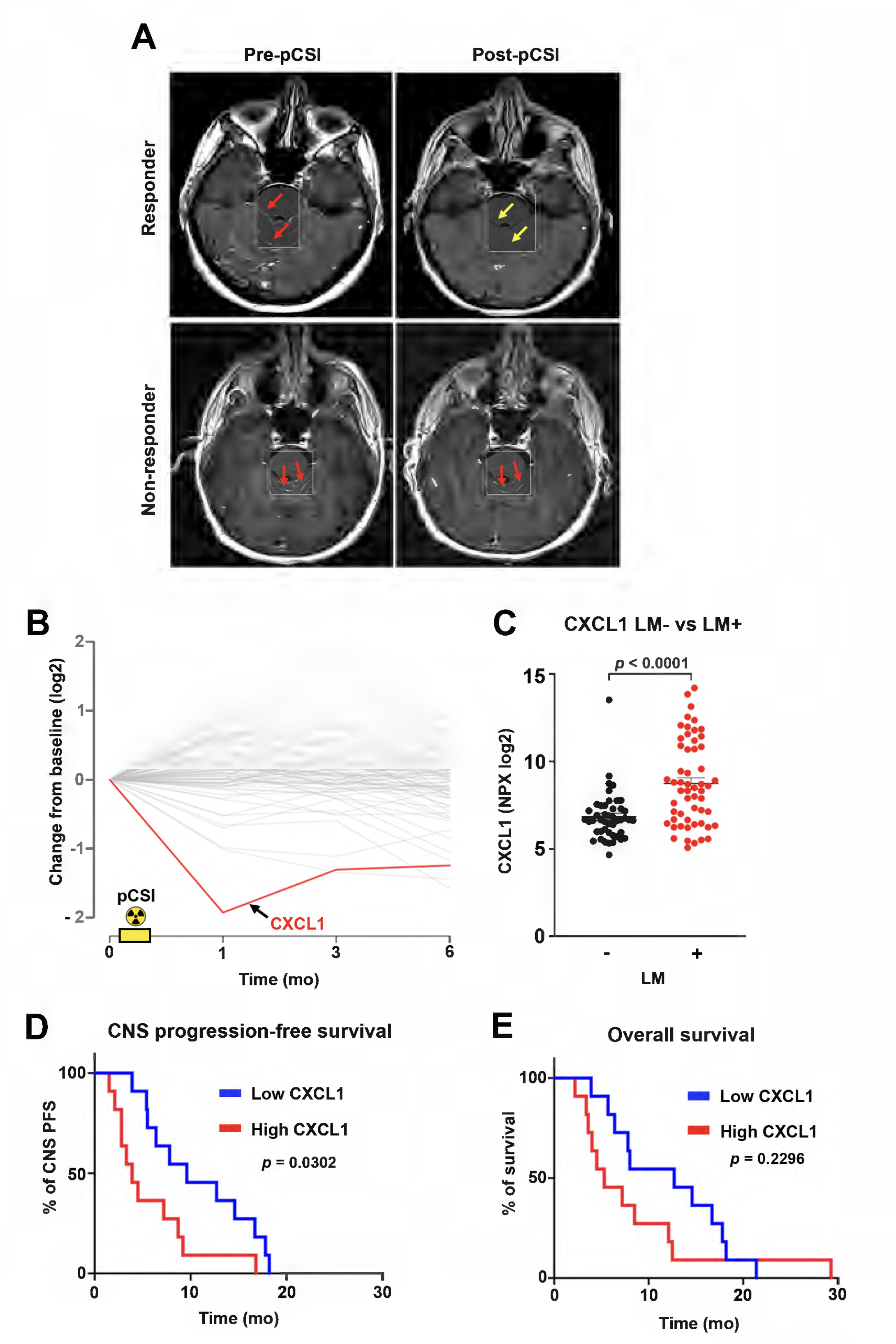
Leptomeningeal metastasis (LM) increases CXCL1 in the cerebrospinal fluid (CSF). (A) Representative axial T1-weighted post-gadolinium contrast enhancement magnetic resonance imaging (MRI) of a responder (upper panel) and non-responder (lower panel) LM patients before and after proton craniospinal irradiation (pCSI) treatment. Sites of radiographically-apparent LM (red arrows) were detectable in non-responder LM patients pre- and post-pCSI, while this was not the case in the responders, where the tumor lesions were resolved due to pCSI (yellow arrows). (B) Dynamics of 92 inflammation-related proteins measured in the CSF from 13 LM patients before and after pCSI using Olink targeted proteomics. Zero (0) time point represents baseline collection prior to pCSI. NPX = Normalized protein expression. For absolute CXCL1 quantification, see Fig.S1B. (C) Dot plot illustrating levels of CXCL1 in the CSF of patients with or without LM, LM+ and LM-, respectively measured using Olink. Data represent mean ± SEM. Unpaired *t*-test. LM-n = 46; LM+ n = 56. *p* < 0.0001. For absolute CXCL1 quantification, see Fig.S1C. (D) Kaplan-Meier curve comparing the central nervous system (CNS) progression-free survival (PFS) of pCSI treated patients stratified based on CXCL1 levels at baseline into low (below median) or high (above median) groups. CXCL1 low n = 11; CXCL1 high n = 11. *p* = 0.0302. (E) Kaplan-Meier curve comparing the overall survival of pCSI treated patients with low or high CXCL1 levels at baseline. CXCL1 low n = 11; CXCL1 high n = 11. *p* = 0.2296.

### Preclinical LM-CSI mouse models recapitulate human disease

To define the role of Cxcl1 in LM growth, we set out to develop an LM-CSI mouse model that faithfully replicates clinical findings observed in human disease. To model LM, we used our established syngeneic LM mouse models of Lewis lung carcinoma (LLC-LeptoM; derived from parental cells of non-small cell lung carcinoma) and triple negative breast cancer (4T1-LeptoM). These cells represent a subpopulation of parental cells generated through iterative *in vivo* selection and readily enter and grow within the leptomeningeal compartment. These cells reliably disseminate along the neuro-axis and mice develop key pathological features and neurological symptoms observed in LM patients (*3, 6, 7*). To model CSI, we tested different time points after intracisternal inoculation of the cancer cells, as well as a variety of irradiation protocols. We first applied a total dose of 10 Gy fractionated over five consecutive days, 2 Gy per fraction (2 Gy × 5). This treatment protocol is commonly used in animal models to control primary brain tumors and CNS metastases, applied in settings of either whole brain irradiation (WBI) or CSI (*15, 16*). Mice received CSI on either day 3, 5, or 7 after inoculation. Of these temporal interventions, initiating CSI on day 7 resulted in a significant reduction of tumor burden compared to untreated sham controls (SH) (fig. S2, A to C); however, CSI with this fractionation regimen did not confer a survival benefit (fig. S2, A to C). We concluded that day 7 was suitable timing to initiate CSI, but this radiation dosage was not sufficient to generate a faithful clinical CSI model. To test this assumption, when we treated the mice on day 7 with 10 Gy CSI given as a single fraction, we obtained both significant reduction in the tumor burden and survival benefit (fig. S2D). We reasoned that five fractions of 2 Gy (total 10 Gy) will achieve a biologically effective dose (BED) of cancer cell death corresponding to 12 Gy, when using the linear-quadratic (LQ) model assuming an α/β ratio of 10 (BED_10_) (*17*), while a single dose of 10 Gy achieves BED_10_ of 20 Gy. On trial, patients were treated with 30 Gy given in 10 fractions (3 Gy × 10), achieving a BED_10_ of 39 Gy. Applying an identical treatment protocol (3 Gy × 10) was not feasible in our LM mouse model due to the narrow time window from inoculation until humane endpoint, corresponding to an average of 14 and 21 days in 4T1-LeptoM and LLC-LeptoM models, respectively (*3, 6, 7*). We therefore applied a hypofractionation treatment protocol: total radiation dose 20 Gy given in 5 fractions (4 Gy × 5; achieves BED_10_ of 28 Gy), often used for WBI for patients with LM (*18, 19*); and adapted it to deliver CSI in mice. This CSI regimen resulted in a significant reduction in tumor burden and conferred survival benefit, in both LLC-LeptoM (Fig. 2, A to C) and 4T1-LeptoM models (Fig. 2, D to F).

**Figure 2:**
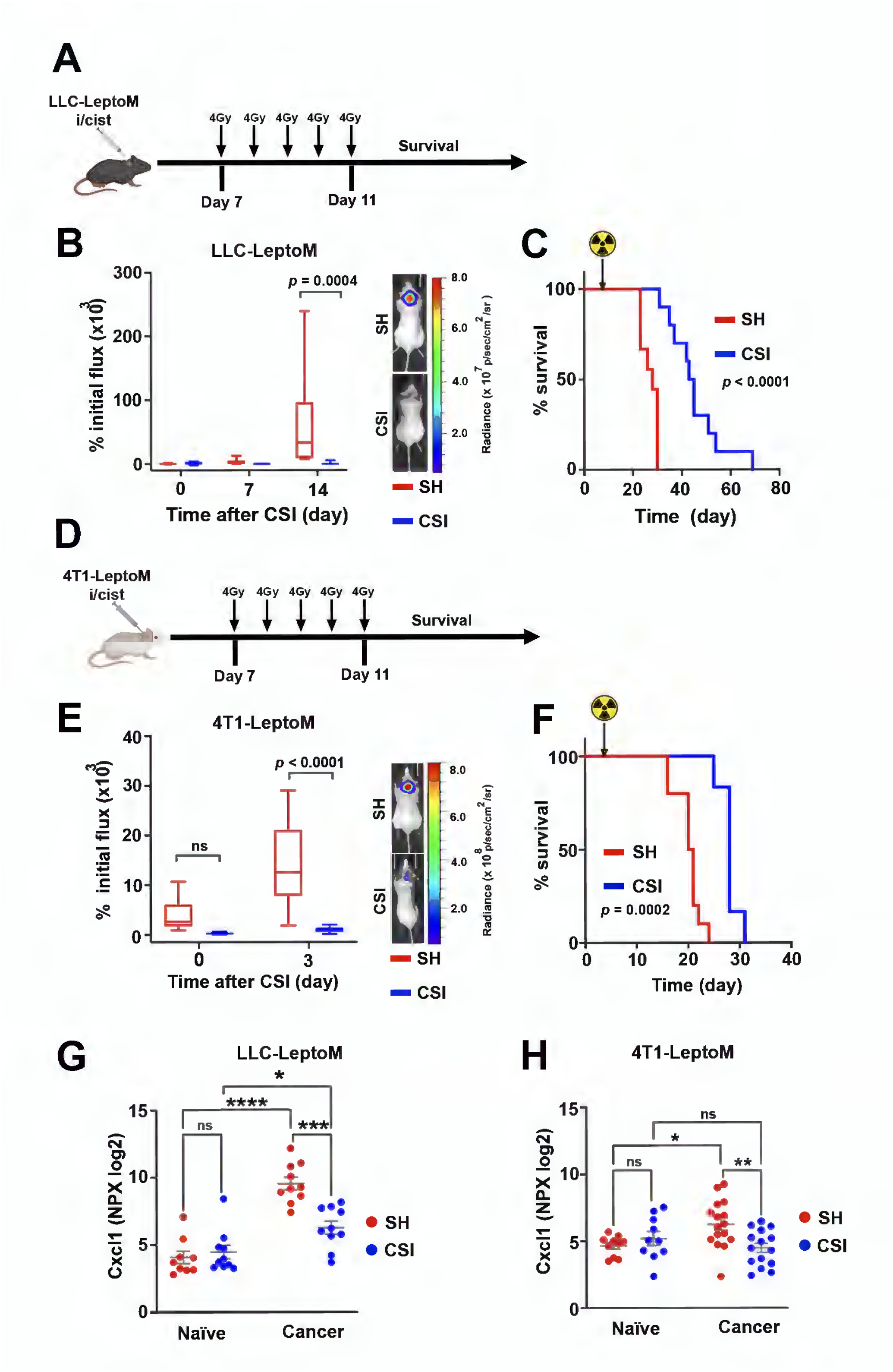
A mouse CSI model recapitulates CXCL1 findings in LM patients. (A) CSI protocol scheme. LLC-LeptoM model is illustrated. i/cist = intracisternal. Gy = Gray. (B) Left: Tumor growth quantified by bioluminescence imaging (BLI). Right: Representative BLI images on day 14 after completion of CSI. Zero (0) = last CSI fraction. Data represent mean ± SEM. Two-way ANOVA with Bonferroni correction. Sham controls (SH) n = 9; CSI n = 10. *p* = 0.0004. (C) Kaplan-Meier survival curve of mice in (**B**). *p* < 0.0001. (D) Scheme for CSI protocol used for 4T1-LeptoM model. (E) Left: Tumor growth quantified by BLI. Right: Representative BLI images on day 3 after completion of CSI. Data represent mean ± SEM. Two-way ANOVA with Bonferroni correction. SH n = 10; CSI n = 8. *P* < 0.0001; ns = not significant. (F) Kaplan-Meier survival curve of mice in (**E**). *p* = 0.0002. (G) Dot plot showing the Cxcl1 levels in mouse CSF from naïve (i/cist injected with PBS) and LLC-LeptoM inoculated mice collected 2 weeks after CSI and measured using the O-link platform. Data represent mean ± SEM. Two-way ANOVA with Tukey’s multiple comparisons test. N = 9 to 10 per group. \**p* = 0.0498, \*\*\**p* = 0.0001, \*\*\*\**p* < 0.0001; ns = not significant. (H) Dot plot showing the Cxcl1 levels in mouse CSF from naïve and 4T1-LeptoM inoculated mice collected 1 week after CSI and measured using the Olink platform. Data represent mean ± SEM analyzed using two-way ANOVA with Tukey’s multiple comparisons test. Naïve n = 10; cancer bearing mice n = 15-16. \**p* = 0.0438, \*\**p* = 0.0081; ns = not significant.

Although both CSI and WBI resulted in comparable control of leptomeningeal tumor growth and survival benefit in our models, WBI treated mice scored higher rates for recumbency and paralysis (60%) compared to those treated with CSI (40%) (fig. S2E). In addition, CSI was superior to spinal irradiation only (SI) (fig. S2F). In our model we used photon (x-ray) irradiation, known to have an exit dose resulting in toxicity in the visceral organs. Our data from irradiated cancer-free animals showed that irradiated animals, regardless of treatment modality, demonstrated slower weight gain compared to SH controls. This trend was significant in the case of cranial irradiation, but not in the case of SI three weeks after completion of treatment (fig. S2G).

With these tools in hand, we next turned to Cxcl1. We measured Cxcl1 in mouse CSF, and found that LM significantly increased CSF Cxcl1 levels, and this elevation was substantially reduced post-CSI, in both mouse models (Fig. 2, G and H). Moreover, CSI alone did not impact the levels Cxcl1 in the CSF of naïve (cancer-free) mice (Fig. 2, G and H). Together, these data demonstrate that our LM-CSI mouse models recapitulate the findings observed in LM patients treated with pCSI, and are suitable for investigating the role of Cxcl1 in the LM setting.

### Multiple cells produce Cxcl1 within the LM microenvironment

To identify the source(s) of increased Cxcl1 within the LM microenvironment, we examined brains and surrounding leptomeninges from mice inoculated with 4T1-LeptoM or LLC-LeptoM cells by immunofluorescence. In both models, we found an increase in the number of Cxcl1-expressing (Cxcl1+) cells in cancer-bearing animals. However, the Cxcl1 expression pattern was distinct between the models (Fig. 3, A and B, and fig. S3, A and B). In the LLC-LeptoM model, hosted in C57Bl6 mice, Cxcl1 expression was abundant, and was observed throughout the leptomeninges (Fig. 3A). In these mice, Cxcl1 was predominantly expressed by macrophage (Iba1+ cells) (Fig. 3, A and C) and Icam1+ leptomeningeal fibroblasts (*20*) (Fig. 3D). Cxcl1 expression was also observed in the choroid plexus, in Iba1+ macrophages and the epithelial cell layer (Fig. 3E). In the 4T1-LeptoM model, hosted in Balb/c mice, Cxcl1 expression appeared minimal in the leptomeninges and displayed a petechial pattern confined to the cancer cells, with no apparent detection in Iba1+ cells (fig. S3, A to C).

**Figure 3:**
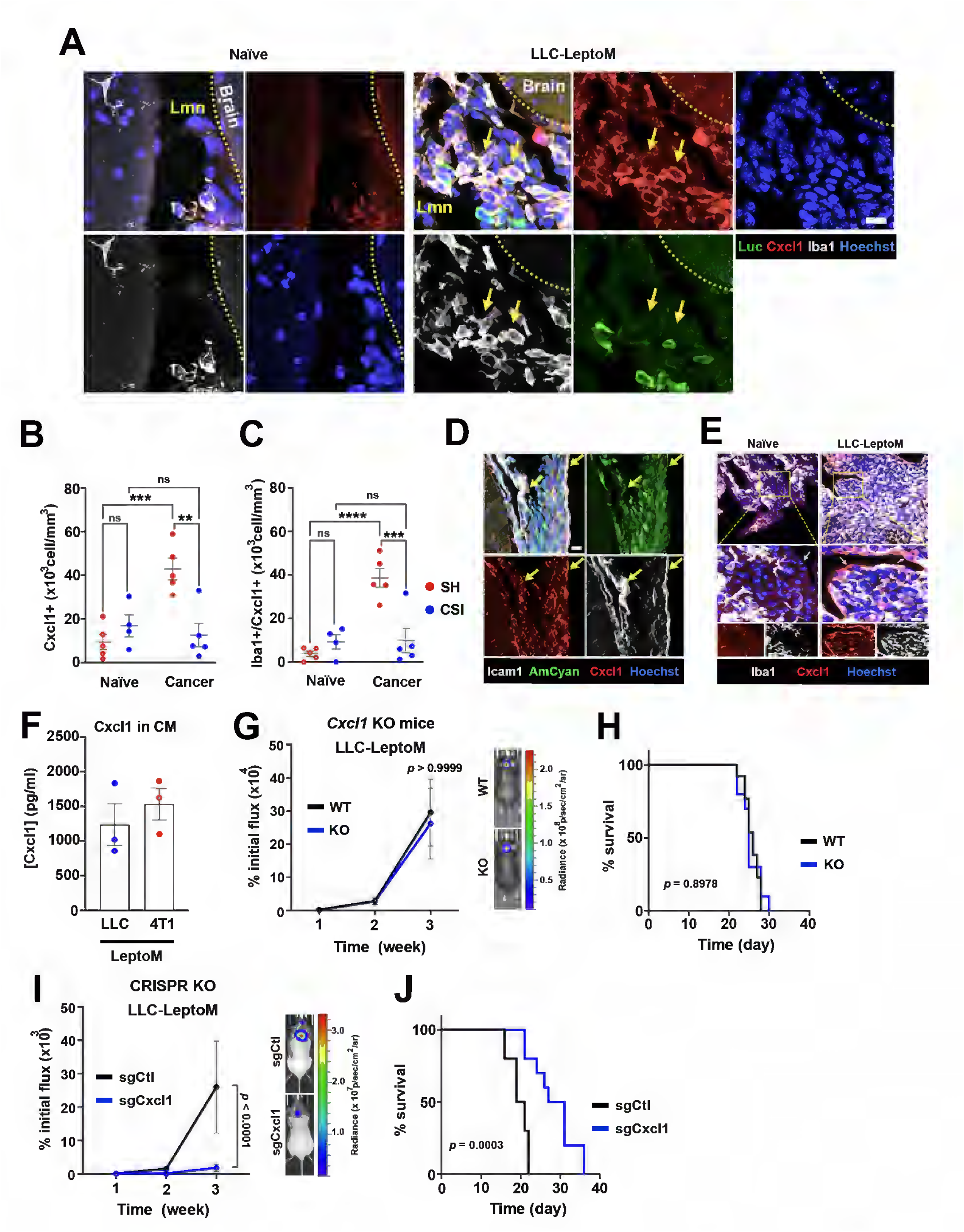
Cancer cells secrete Cxcl1 to support their leptomeningeal growth. (A) Confocal images illustrating Cxcl1 expression in leptomeninges (Lmn) of naïve and LLC-LeptoM inoculated mice 3 weeks after inoculation of cancer cells. Green: luciferase (Luc) expressed by cancer cells; Red: Cxcl1; White: Iba1, expressed by macrophages; Blue: Hoechst, nuclear counter stain. Scale bar = 10 μm. (B) Dot plot quantification of Cxcl1 expressing cells (Cxcl1+) in leptomeninges of naïve and LLC-LeptoM inoculated mice 2 weeks after CSI. Data represent mean ± SEM Two-way ANOVA with Tukey’s multiple comparisons test. n = 4-5 per group. \*\**p* = 0.0016, \*\*\**p* = 0.0006; ns = not significant. (C) Dot plot showing quantitation of Iba1+ cells co-expressing Cxcl1 (Iba1+/Cxcl1+) in leptomeninges of naïve and LLC-LeptoM inoculated mice 2 weeks after CSI. Data represent mean ± SEM. Two-way ANOVA with Tukey’s multiple comparisons test. n = 4-5 per group. \*\*\**p* = 0.0006, \*\*\*\**p* < 0.0001; ns = not significant. (D) Confocal images showing Cxcl1 (red) co-expression by leptomeningeal fibroblasts (Icam1+ cells; white) in cancer-bearing animals, indicated with yellow arrows. Tissue collected 3 weeks after inoculation of LLC-LeptoM (labeled with AmCyan, green); Blue: Hoechst, nuclear counter stain. Scale bar = 20 μm. (E) Confocal images showing Cxcl1 (red) in the choroid plexus of cancer-bearing animals. Cxcl1 co-expression was observed in Iba1-expressing cells (white), as well as in the epithelial cell layer (white arrows). Tissue collected 3 weeks after inoculation of LLC-LeptoM. Blue: Hoechst, nuclear counter stain. Scale bar = 50 μm (overview image) and 10 μm (close up image). (F) Histogram of Cxcl1 levels in conditioned media (CM) collected from LLC-LeptoM and 4T1-LeptoM grown in serum-free media for 24 h. n = 3 independent experiments. (G) Tumor growth in wildtype mice (WT) vs Cxcl1 knockout (KO) mice. Left: Tumor growth quantified by BLI. Right: Representative BLI images 3 weeks after inoculation of LLC-LeptoM cells. Data represent mean ± SEM. Two-way ANOVA with Bonferroni correction. WT n = 13; KO n = 10. *p* > 0.9999. (H) Kaplan-Meier survival curve of mice in (**G**). *p* = 0.8978. (I) Tumor growth of Cxcl1 KO LLC-LeptoM cells (sgCxcl1) vs control cells (sgCtl). Left: Tumor growth quantified by BLI. Right: Representative BLI images 3 weeks after inoculation of cancer cells. Data represent mean ± SEM. Two-way ANOVA with Bonferroni correction. sgCtl n = 10; sgCxcl1 n = 10. *p* < 0.0001. (J) Kaplan-Meier survival curve of mice in (**I**). *p* = 0.0003.

Because capture of a secretory protein by immunofluorescence is often challenging unless its expression is abundant, we hypothesized that immunofluorescence may have failed to visualize Cxcl1 expression in the cancer cells in the LLC-LeptoM model due to the relative abundance in surrounding Iba1+ cells. Using ELISA, we measured the levels of Cxcl1 in conditioned media (CM) obtained 24 h after growing LLC-LeptoM and 4T1-LeptoM cells in serum-free media. We found that both LeptoM models secreted Cxcl1 (Fig. 3F), and irradiation did not alter cancer cell Cxcl1 production (fig. S3D). Together, these results indicate that both cancer and host cells contribute to elevated Cxcl1 levels detected in the CSF after LM.

### Cancer cell-derived Cxcl1 supports leptomeningeal cancer cell growth

To address the functional role of Cxcl1 in LM growth, we employed a bi-directional genetic approach to interrupt Cxcl1 production from the host and the cancer cells. To target host-generated Cxcl1, *Cxcl1*-/-mice (*21*) and their +/+ littermates (WT) were inoculated with LLC-LeptoM. We did not observe differences in tumor growth or overall survival between the two genotypes (Fig. 3, G and H). To target cancer cell-generated Cxcl1, we knocked out *Cxcl1* with CRISPR/cas9 using two independent single guide (sg) RNA against *Cxcl1* and one non-targeting control guide (LacZ). Although Cxcl1 production was successfully interrupted as measured by ELISA (fig. S3E), its loss did not impact cancer cell growth *in vitro* (fig. S3F). In contrast, *in vivo*, cancer cell loss of Cxcl1 suppressed cancer cell growth in the leptomeninges and improved survival (Fig. 3, I and J, and fig. S3G). Importantly, interrupting cancer cell Cxcl1 expression did not suppress tumor growth in the lung (the orthotopic primary site), or subcutaneous tissue (fig. S3, H and I). In addition, treatment with recombinant Cxcl1 *in vitro* did not alter cancer cell growth (fig. S3J). Taken together, these results demonstrate that cancer cell-derived Cxcl1 supports cancer cell growth in the leptomeningeal compartment, and this effect is specific to the leptomeningeal microenvironment.

### Cancer cell Cxcr2 expression confers a leptomeningeal growth advantage

As the effect of Cxcl1 on LM growth was site-specific and not captured *in vitro*, we next explored this signaling axis within the leptomeninges. In mice, Cxcl1 primarily binds the C-X-C motif chemokine receptor 2 (Cxcr2) to induce its biological actions, including immune cell trafficking (*22*). To characterize Cxcr2 expression within the LM microenvironment, we began by assaying Cxcr2 expression in leptomeningeal CD45+ cells in LLC-LeptoM and 4T1-LeptoM models using flow cytometry. LM is well known to provoke a robust inflammatory response (*8*). Accordingly, we found that the leptomeninges in the setting of LM contained a host of immune cells (fig. S4, A and B). Each model generated a distinct array of immune cell responses to LM with Cxcr2 expression patterns specific to the mouse genetic background (fig. S4, A to D), as observed in other models (*23–25*). To address the role of Cxcr2 expressed by immune cells in LM growth, we leveraged a hematopoietic-specific Cre-recombinase to delete *Cxcr2* in cells originating from hematopoietic progenitors, including myeloid and lymphoid cells (*26–28*). Interestingly, deletion of Cxcr2 in this population promoted leptomeningeal cancer cell growth (fig. S4E), suggesting that immune cells do not mediate the pro-tumorigenic effect of Cxcl1. We then turned our attention to cancer cell Cxcr2 expression. Evaluation of Cxcr2 expression in LeptoM cells by flow cytometry revealed that these cells were heterogenous with regard to Cxcr2 expression. LLC-LeptoM and 4T1-LeptoM cells contained both Cxcr2-negative (Cxcr2-) and relatively rare (0.02-0.5%) Cxcr2-expressing (Cxcr2+) cells (Fig 4, A and B, fig. S4F). Interestingly, although the proportion of Cxcr2+ cells remains stable *in vitro* (fig. S4F), after inoculation and growth within the leptomeningeal compartment, the proportion of Cxcr2+ cells sharply elevated to 35% and 20% of the total cancer cells in LLC-LeptoM and 4T1-LeptoM models, respectively (Fig 4, A and B, fig. S4G). To examine the functional relevance of cancer cell Cxcr2 expression in the context of the leptomeninges, we used flow cytometry to sort the LeptoM cell lines into Cxcr2- and Cxcr2+ subpopulations, and instilled these into the leptomeninges. Cxcr2+ cancer cells demonstrated a growth advantage within the leptomeninges (Fig 4, C and D). To understand the transcriptional programming of these subpopulations, Cxcr2- and Cxcr2+ LeptoM cells were isolated and subjected to bulk RNA-sequencing (RNA-seq). Principal component analysis (PCA) demonstrated distinct clustering of Cxcr2- and Cxcr2+ cells in both LM models (Fig. 4E), indicating substantial global transcriptional differences between these subpopulations. In LLC-LeptoM cells, we found a total of 156 differentially expressed genes (DEGs) between Cxcr2- and Cxcr2 + cells, of which 149 and 7 genes were up- and downregulated, respectively (Fig. 4F). In 4T1-LeptoM cells, DEGs between these two subpopulations were 295, of which 259 and 36 genes were up- and downregulated, respectively (Fig. 4F, Table S1). Moreover, we found 34 upregulated genes in common across both LeptoM cell lines; there was no overlap detected between the downregulated genes (Fig. 4F). Examining the DEG by gene ontology analysis, we uncovered an enrichment of pathways related to cell cycle progression in Cxcr2+ cells in both LeptoM models (Fig. 4G). Taken together, these results demonstrate a selective advantage of Cxcr2 expression in cancer cells linked to their progression within the leptomeningeal compartment, suggesting that signaling through this receptor is likely crucial for promoting LM growth.

**Figure 4:**
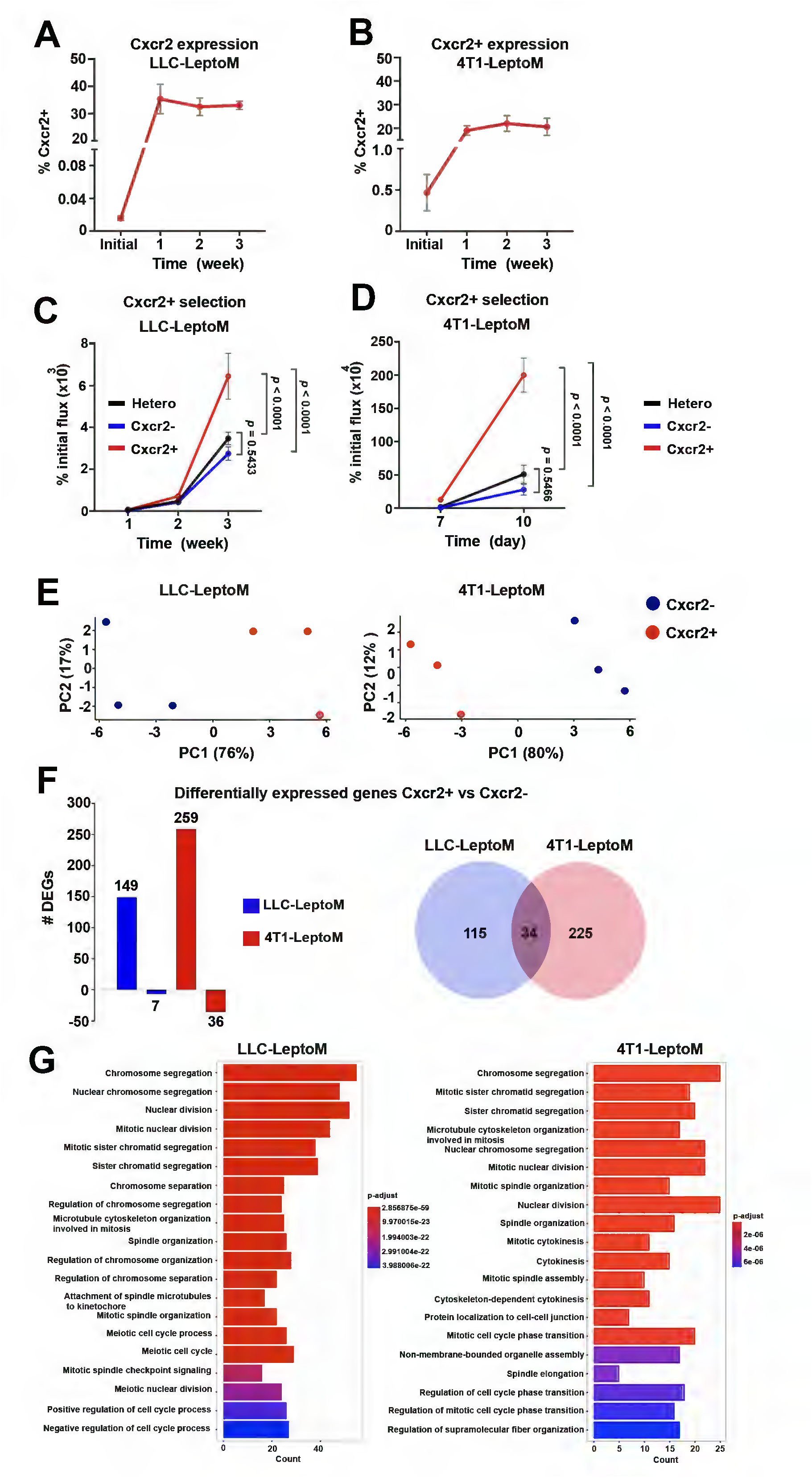
Cancer cell Cxcr2 expression confers a leptomeningeal growth advantage. (A) Percentage of Cxcr2 expressing (Cxcr2+) LLC-LeptoM cells before (initial) and after i/cist inoculation as quantified by flow cytometry. Initial: n = 4 independent experiments. i/cist: n = 5 per time point. (B) Percentage of Cxcr2+ 4T1-LeptoM cells before (initial) and after i/cist inoculation as quantified by flow cytometry. Initial: n = 4 independent experiments. i/cist: n = 5 per time point. (C) Tumor growth of selected Cxcr2+, Cxcr2 negative (Cxcr2-) and the original heterogeneous (Hetero) LLC-LeptoM cells as quantified by BLI. Data represent mean ± SEM analyzed using two-way ANOVA with Bonferroni correction. Hetero: n = 9; Cxcr2-n = 8; Cxcr2+ n = 8. Hetero vs Cxcr2-*p* = 0.5433; Hetero vs Cxcr2+ *p* <0.0001; Cxcr2-vs Cxcr2+. *p* < 0.0001. (D) Tumor growth of selected Cxcr2+, Cxcr2 negative (Cxcr2-) and the original heterogeneous (Hetero) 4T1-LeptoM cells as quantified by BLI. Data represent mean ± SEM analyzed using two-way ANOVA with Bonferroni correction. Hetero: n = 9; Cxcr2-n = 8; Cxcr2+ n = 8. Hetero vs Cxcr2-*p* = 0.5466; Hetero vs Cxcr2+ *p* <0.0001; Cxcr2-vs Cxcr2+. *p* < 0.0001. (E) Principal component analysis (PCA) plot showing distinct transcriptional clustering of Cxcr2- and Cxcr2+ cell populations in both LLC-LeptoM and 4T1-LeptoM cell lines. (F) Left: Histogram of differentially expressed genes (DEGs) between Cxcr2- and Cxcr2+ cells in LLC-LeptoM and 4T1-LeptoM cell lines. Right: Venn diagram illustrating unique and shared DEGs between LeptoM cells lines. (G) Gene ontology analysis of top 20 significantly enriched pathways in Cxcr2+ cells of LLC-LeptoM (left) and 4T1-LeptoM (right).

### Locoregional interruption of Cxcr2 signaling suppresses LM growth

Because expression of the Cxcl1-Cxcr2 axis supported LM growth, we speculated that interruption of Cxcr2 signaling might suppress LM growth. Accordingly, we interfered with Cxcr2 signaling with the selective antagonist SB265610 (SB) via two interventional routes: systemic intraperitoneal (i/p) delivery and local intracisternal delivery specifically to the CSF (i/cist), using the LLC-LeptoM model. For the i/p treatment, LeptoM cells were i/cist inoculated and treatments with SB or vehicle (Veh; 10% DMSO) were started immediately and maintained daily for 3 weeks, single injection a day (fig S5A). Using this route of administration, we neither observed suppression of cancer growth nor survival benefit (fig S5, B and C). For i/cist treatment, LeptoM cells were co-inoculated with SB or vehicle, and treatments were maintained i/cist for 2 weeks, with an interval of 48 h and 72 h during the first and second week, respectively (Fig. 5A). Using this route of administration, interference with Cxcr2 substantially suppressed LM growth and conferred survival benefit to the treated animals (Fig. 5, B and C). Notably, we did not observe overt toxicity from the i/cist treatment, as judged by the overall weight gain on the last treatment day (fig S5D). To examine pharmacological interruption of Cxcr2 signaling in a more clinically relevant context, we initiated i/cist treatment of vehicle or SB 7 days after inoculation of LLC-LeptoM cells, with or without a lower dose of CSI (total 10 Gy; 2 Gy × 5; BED 12 Gy) (Fig. 5D). This CSI dose does not confer survival benefit alone, as demonstrated above (fig. S2C). While each of these treatments suppressed cancer cell growth, combination of SB with CSI was the only intervention that conferred survival benefit (Fig. 5, E and F). Collectively, these results demonstrate that locoregional interruption of Cxcr2 signaling inhibits cancer cell growth within the LM microenvironment, and combination with CSI potentiates the efficacy of Cxcr2 inhibition.

**Figure 5:**
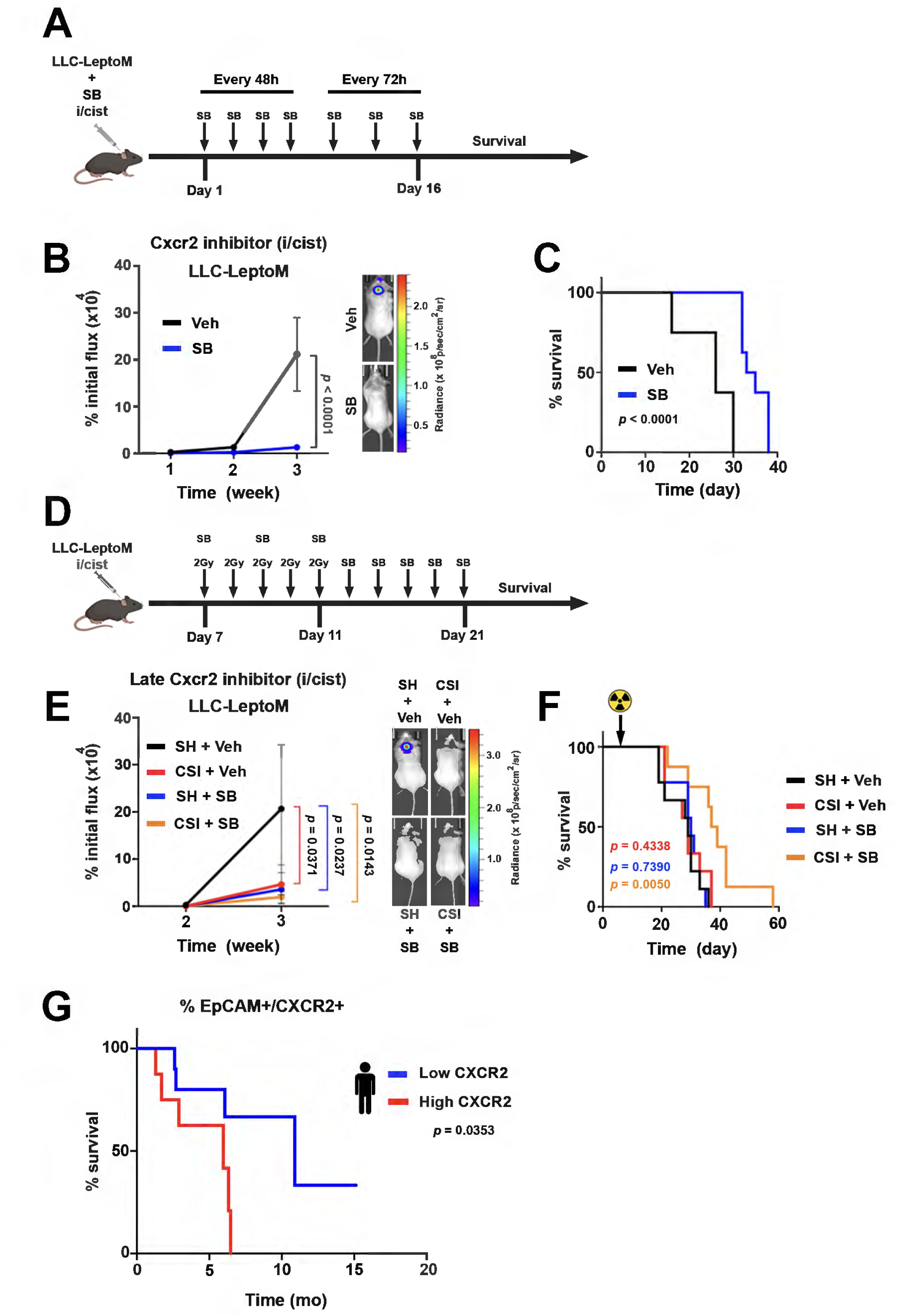
Pharmacologic interruption of Cxcr2 inhibits LM growth and promotes survival. (A) Experimental treatment scheme illustrating i/cist treatment with Cxcr2 inhibitor SB265610 (SB) or vehicle (Veh) in LLC-LeptoM model. Treatments were co-injected with cancer cells and maintained for 2 weeks. (B) Left: Tumor growth as quantified by BLI. Right: Representative BLI images 3 weeks after inoculation of cancer cells. Data represent mean ± SEM. Two-way ANOVA with Bonferroni correction. Veh n = 8; SB n = 8. *p* < 0.0001. (C) Kaplan-Meier survival curve of mice in (**B**). *p* < 0.0001. (D) Scheme showing the paradigm of late treatment with Cxcr2 inhibitor SB with or without CSI (10 Gy delivered as 2 Gy × 5 fractions). Treatments initiated one week after inoculation of cancer cells and continued for 2 weeks. (E) Left: Tumor growth quantified by BLI. Right: Representative BLI images 3 weeks after inoculation of cancer cells. Data represent mean ± SEM. Two-way ANOVA with Bonferroni correction. SH + Veh n = 9; CSI + Veh n = 9; SH + SB n = 9; CSI + SB n = 8. SH + Veh vs CSI + Veh *p* = 0.0371; SH + Veh vs SH + SB *p* = 0.0237; SH + Veh vs CSI + SB *p* = 0.0143. (F) Kaplan-Meier survival curve of mice in (**E**). SH + Veh vs CSI + Veh *p* = 0. 4338; SH + Veh vs SH + SB *p* = 0.7390; SH + Veh vs CSI + SB *p* = 0.0050. (G) Kaplan-Meier curve comparing the overall survival of LM patients stratified based on the percentage of CXCR2 expression in cancer cells (EpCAM+/CXCR2+) in CSF prior to pCSI. CXCR2 low (less than 1%) n = 10; CXCR2 high (more than 1%) n = 8. *p* = 0.0353. mo = month.

### Cancer cell CXCR2 expression in LM patients portends poor response to pCSI

Finally, to gain a translational insight into the association of CXCR2 with LM progression, we asked if the findings observed in our mouse models are also mirrored in patients. We evaluated CXCR2 expression in LM originating from a variety of solid primary carcinomas in CSF collected prior to pCSI (Table S2). Using flow cytometry, we analyzed Cxcr2 expression in cancer cells, identified by expression of epithelial cell adhesion molecule (EpCAM), after exclusion of CD45+ immune cells. We found that higher proportion of CXCR2 expression in cancer cells (EpCAM+/CXCR2+) prior to pCSI correlated with worse overall survival after pCSI (Fig. 5G).

## Discussion

Patients with leptomeningeal metastases have long been excluded from the vast majority of clinical trials (*29*). As a result, formal clinical and translational investigation of novel treatments for LM remains rare (*10*). However, unlike other central nervous system metastases, LM offers the opportunity for serial collection of the metastatic site in human disease. Collection of CSF is integral to diagnosis and management of LM (*30*). We have leveraged these collections in the context of a clinical trial to dissect the biology of treatment response and resistance.

Current clinical metrics for monitoring of LM treatment response rely on MRI and CSF cytology to assess LM progression (*30*). Enhancement in MRI over the surface of the brain or spinal cord has been useful to visualize LM plaques, however, imaging has significant limitations, including the procedural burden for the patient and lower sensitivity and specificity (*31–33*). While CSF cytology represents the “gold standard” for LM diagnosis, the test suffers from notoriously low sensitivity, necessitating serial CSF collections (*31, 33*). Our findings linking CSF CXCL1 to the presence of LM and response to pCSI suggest that CSF CXCL1 measurement has the potential to serve as a robust biomarker of treatment response, facilitated via routine liquid biopsy.

CSF biomarkers may have the potential to aid in predicting and monitoring response to pCSI. Previous work has employed CSF circulating tumor cell (CTC) counts (*34*). Unfortunately, current CTC detection systems are not universal. Because these techniques are largely based on the detection of EpCAM, they are limited to cancer cells of epithelial origin, and have a limited dynamic range (an upper limit of detection of 200 CTC / 3 ml) (*35*). In this study, targeted proteomic screen of CSF identified the chemokine CXCL1 as a potential biomarker of LM. CSF CXCL1 levels correlate with both presence of cancer cells in the leptomeningeal compartment and response to pCSI in patients. Measurement of a protein such as CXCL1 offers many advantages over cell-based biomarkers. Whereas CSF CTCs require specialized collection tubes and instrumentation, CXCL1 levels can be accurately assayed in smaller volumes of CSF with simple tests, including ELISA.

Recently, pCSI has emerged as a promising treatment modality prolonging the survival of LM patients. The technique enables precise delivery of radiation to the entire neuro-axis overcoming many of the toxicities associated with conventional photon irradiation (*11, 36*). However, toxicities, notably fatigue and memory loss, remain (*37*), and the treatment requires substantial patient commitment. Because response to pCSI varies substantially between patients, identification of patients likely to derive benefit from this therapy represents a major unmet need.

Beyond biomarker discovery, our work has uncovered a novel signaling axis in LM. Previously, CXCL1 has been observed to play a role in cancer growth, dissemination, and resistance to therapies in multiple organs, including the brain (*38, 39*). Here, we find that cancer cell-derived CXCL1 is crucial for cancer cell growth within the leptomeningeal compartment, uncovering a unique site-specific signaling axis mediating cancer cell growth within the leptomeninges (*6, 7*). In mice, CXCL1 induces its biological actions primarily through binding to the CXCR2 receptor (*22*). Although other chemokines can also signal through this receptor (*22, 40–42*), we have centered this work on the CXCL1-CXCR2 axis, as CXCL1 was uncovered in an unbiased fashion and robustly correlated with outcomes in human disease. CXCL1-CXCR2 signaling has been shown to play pro-tumorigenic and metastatic roles by recruiting immune suppressor cells that can dampen the action of anti-tumor immune effectors or by precipitating secretion of factors supporting cancer cell growth and dissemination (*38, 41–43*). In contrast, in the leptomeningeal space, we find that signaling through CXCR2 in immune cells appeared to exert an anti-tumor effect. Genetic loss of CXCR2 expression in immune cells promoted cancer growth. Conversely, CXCR2 expression in cancer cells conferred a growth advantage within the leptomeningeal compartment, underlining the unique pro-tumor signaling that occurs within this space (*6, 8*).

Our work has captured dynamic changes within cancer cells as they inhabit the leptomeninges. CXCR2-expressing cancer cells demonstrate a clear growth advantage within this space. Molecular profiling of CXCR2-expressing cancer cells further revealed enrichment of pathways linked to cell cycle progression, suggesting that CXCR2 expression is important for cancer cell propagation within the leptomeningeal compartment. In support of this, many of the genes we found enriched in the CXCR2-expressing cancer cell subpopulation have previously been associated with cancer stem cells in multiple cancer types, including breast, lung, and colorectal cancers (*44–46*). Indeed, we found that locoregional interruption of CXCR2 was sufficient to suppress LM growth. Inhibition of cancer cell growth and metastasis via interference with CXCR2 has previously been demonstrated in several cancer types (*47–49*), but has not been reported in the context of LM.

We also show that CXCR2 inhibition is synergistic in combination with CSI. This suggests that addition of CXCR2 inhibition with pCSI may permit reduction of radiation dose, and hence minimizing the neurotoxicity resulting from cranial irradiation, and improving the quality of life (*50*). In addition, CXCR2 inhibition might be valuable for targeting LM growth in cancer types or in patients for whom radiation is not a treatment option (*49, 51*).

Overall, we have demonstrated that CSF CXCL1 correlates with resistance to pCSI and may serve as a biomarker to predict and monitor response to treatment. In addition, we find that the CXCL1-CXCR2 signaling axis represents an actionable therapeutic target to interrupt LM growth. Combination of pCSI with CXCR2 inhibition represents a promising and actionable therapeutic strategy to suppress progression of this deadly complication of cancer (*22, 52*).

## Acknowledgments

We wish to express our deep gratitude to the patients and families that enrolled in NCT03520504 and NCT04343573, as well as those outside these trials that donated clinical samples enabling this research. We also would like to thank the Radiation Facility, Molecular Cytology, Flow Cytometry, and Integrated Genomics Operation Core Facilities at MSCC. These Core Facilities are funded, in part, by the NCI Cancer Center Support Grant (CCSG, P30 CA08748), Cycle for Survival, and the Marie-Josée and Henry R. Kravis Center for Molecular Oncology.

## Author contributions

A.B. supervised the study. A.M.O., J.R., and A.B. conceived and designed the study. A.B. acquired funding. A.M.O. and AB. wrote the manuscript. J.R. and X.T. generated metastatic cell lines. A.M.O., B.M., A.R.N., D.G., H.W., and M.J.L. performed the experiments. A.M.O., D.G., and S.P-O performed *in vitro* experiments. A.M.O., A.R.N., and D.G. performed the ELISA. B.M., A.R.N., D.G., and J.R. performed flow cytometry analysis and assisted with designing CRISPR knockout experiments. J.S., performed RNA extraction and analysis of RNA-sequencing data. A.J.D. and R.L.L. contributed to the Cxcr2 knockout experiment. N.A.W performed computational analysis of the Olink data. K.C., M.F., J.T.Y., and AB. annotated the patient material and clinical data. All authors approved the final manuscript.

## Material and data availability

RNA-seq datasets were deposited in the NCBI Gene Expression Omnibus (GEO) under the accession numbers GSE283519. Commercially available materials can be obtained from vendors, details are available in material and methods. Materials generated in this study are available from the corresponding author upon signing the MSKCC Material Transfer Agreement. Human samples used in this study are limited biological resource, not available for further distribution.

## Supplementary materials and methods

### Study design

This study aimed to define the molecular basis of leptomeningeal metastasis resistance to pCSI. We collected cell-free fractions of CSF from cancer patients with and without LM, and those envisioned to receive pCSI before and throughout the course of treatment. This cell-free CSF was subjected to targeted proteomic screens using proximity ligation assay to identify possible targets associated with LM growth and resistance to treatment. The identified targets were further validated using ELISA. We also developed syngeneic LM-CSI models that faithfully recapitulated our observations in patient samples. We leveraged these models to investigate the source of these cytokines, their cellular target(s), and the role of these targets in LM growth. We performed functional studies using both genetic and pharmacological approaches *in vivo* and *in vitro*. We carried out transcriptomic analysis of LM cell subpopulations using bulk RNA sequencing. The sample sizes for studies carried out with human samples were based on available samples, and were therefore limited. The number of animals used in each experiment was based on our previous work using these models: With 10 mice in each group (control vs. knockdown), we have approximately 80% power to detect a 20% difference in defined cell populations between the two groups assuming a standard deviation of 11070 from bioluminescent signal data using a two-sided Wilcoxon rank-sum test at the 0.05 significance level. All animal experiments were performed in at least two independent experiments. For the breast cancer model females were used, and for lung cancer models, both females and males were used at 1:1 ratios. All *in vitro* studies were performed in three independent experiments, where each constituted at least three technical replicates. All histological quantification was performed blinded. Sample sizes and statistical tests are indicated in the figure legends and in the “Statistical analysis” section.

### Human studies

Cerebrospinal fluid (CSF) was collected from patients who provided consent. In the case of pCSI studies, the patients carried a diagnosis of leptomeningeal metastasis (LM) and enrolled in NCT03520504 and NCT04343573. For studies examining CSF from patient with and without LM, CSF was collected from patients undergoing CSF collections for clinical indications. CSF in excess of that required for clinical decision making was banked. For all patients, LM was diagnosed by both radiographic criteria on magnetic resonance imaging (MRI) and positive CSF cytology. Both clinical trials and biospecimen studies were approved by Memorial Sloan Kettering Cancer Center (MSKCC) Institutional Review Board (IRB; protocols: 06-107, 12-245, 13-039, 18-205, 18-505 and 20-117. In the case of pCSI trials, CSF collections were performed starting approximately two weeks prior to the first pCSI session (baseline) and at multiple points after completion of pCSI (∼1-, 3- and 6-months), as per trial protocol. For all patients, CSF was collected either through the lumbar puncture or via Ommaya reservoir, spun at 600 × g for 5 min at 4°C to generate cell-free CSF and cellular fraction. The cell-free CSF was stored at −80°C, and the cellular fractions were cryopreserved using serum-free cryo-preservant (GC Lymphotec, Bambanker BB01) and stored at −80°C until further use.

### Targeted Proteomics

Samples were processed and analyzed essentially as described in (*53*). CSF collected between 2015-2020 was aliquoted and stored at −80°C at MSK Brain Tumor Center CSF Bank. Samples were slowly thawed on ice and 45 μL of CSF was mixed with 5 μL of 10% Triton X-100 (Sigma, T8787) in saline and incubated at room temperature for two hours (final concentration of Triton X-100 was 1%). Samples were then dispensed into 96-well PCR plates and stored at −80°C until further analysis. Relative levels of proteins in two targeted panels were detected using proximity extension assay (Olink Target 96 Inflammation). Protein abundance values are shown in NPX units (*log2* scale). The analytical range for each analyte is available online (www.olink.com).

### Animals

Animal experiments were approved by MSKCC Institutional Animal Care and Use Committee (IACUC) protocol number 18-01-002. The following mouse strains were used: B6 albino mice (B6(Cg)-*Tyr^c-2J^*/J; stock # 000058), BALB/cJ (stock # 000651). Mice for experiments from these two strains were obtained from Jackson laboratory. *Cxcl1* mutant C57BL/6NCrl-*Cxcl1^em1(IMPC)Mbp^*/Mmucd, RRID:MMRRC_046310-UCD, obtained from the Mutant Mouse Resource and Research Center (MMRRC) at University of California Davis (*21*). *Cxcr2* knock-out mice (KO) were generated by crossing *Cxcr2* floxed mice to *Vav-Cre* lines as described previously (*26, 54*). *Cxcl1* mutant and *Cxcr2* KO mice were bred in-house, and the litter genotypes were determined per provider protocols. Mice were used at the age of six to eight weeks. For the breast cancer model, females were used. For the lung cancer model, both females and males were used in approximately 1:1 ratio. Animals were housed in individually ventilated cages in an equal light-dark cycle with *ad libitum* access to food and water.

### Cell lines and cell culture

Mouse Lewis lung carcinoma (LLC) and mouse triple-negative breast cancer 4T1 cell lines were used. Both cell lines express the firefly luciferase and underwent iterative *in vivo* selection to generate their metastatic leptomeningeal derivatives as described previously (*6, 7*), henceforth referred to as LLC-LeptoM and 4T1-LeptoM. In some experiments, we used their derivatives engineered to express the AmCyan or mCherry fluorescent proteins (see figure legends). LLC-LeptoM and 4T1-LeptoM cells were grown in DMEM or RPMI culture media, respectively. Culture media were supplemented with 10% fetal bovine serum (FBS; Omega Scientific #FB-11), 1% penicillin-streptomycin (ThermoScientific #15140-122), and 1 % L-glutamin-derivative GlutaMAX (ThermoScientific #35050-061). Cells were maintained at 37°C and 5% CO_2_, and frequently tested for mycoplasma contamination using MycoAlert™ mycoplasma detection kit (Lonza #LT07-218).

### Conditioned media collection

Conditioned media (CM) from LLC-LeptoM and 4T1-LeptoM cells were generated to evaluate secretion of Cxcl1 produced by either cell line. Cells were plated in 6-well plates at a density of 1.5 × 10^5^ in complete culture media containing 10% FBS, and allowed to settle overnight. Plates were then carefully rinsed three times with phosphate-buffered saline (PBS), and serum-free medium supplemented with 0.5% bovine serum albumin (BSA; Miltenyi #130-091-376) was added. CM were collected 24 h thereafter, spun at 1000 × g for 10 min at 4°C to remove floating live and dead cells and debris, and the supernatants were transferred into new tubes and stored at −80°C until assayed.

### Generation of Cxcl1 knock-out cancer cell lines

Cxcl1 knockout in LLC-LeptoM was performed using CRISPR/Cas9 as previously described (*6*). Lentivirus vectors were designed to deliver control single guide (sg) RNA (*LacZ*; sg*LacZ* ‘TGCGAATACGCCCACGCGAT’), or two independent sgRNAs targeting mouse *Cxcl1* (sg*Cxcl1* ‘CTTGAGGTGAATCCCAGCCA’ and sg*Cxcl1*‘ACTTCGGTTTGGGTGCAGTG’; VectorBuilder). Plasmids were extracted using ZymoPureII plasmid maxi prep kit (Zymo Research Cat #D4203). Virus particles were generated in Lenti-X™ 293T cell line (Clontech #632180) using Lenti-X™ packaging single shots (VSV-G) Clontech #631276. Supernatants were collected, and the virus titer was concentrated using LentiX concentrator (Takara Cat# 631232). LLC-LeptoM were then transduced with lentivirus and selection of the successfully transduced cells was performed using Blasticidin (10 μg/ml; Invivogen #ant-bl-1). Cells were then sorted into 96-well plate using FACSAriaIII (BD Biosciences), single cell per well. The absence of Cxcl1 secretion in successfully expanded clones was screened and quantified with ELISA.

### Cell growth assay

To assess the growth of LLC-LeptoM and 4T1-LeptoM in response to Cxcl1 treatment, cells were plated in a flat-bottom 96-well plate at a density of 2 × 10^3^/well supplied with complete media and allowed to adhere overnight. After washing with PBS, the culture media were switched to serum-free, and cells were treated with either vehicle (PBS) or different concentrations of carrier-free recombinant mouse Cxcl1 (R&D Systems #453-KC/CF) ranging from 0.01 to 1 ng/ml, corresponding the ranges found in the CSF of LM patients (Fig.S1 B and C). Cell growth was evaluated 24 h after treatment using the cell titer aqueous proliferation assay kit (Promega #G3582). To assess the cell growth after Cxcl1 knockout, control and Cxcl1 knockout LLC-LeptoM cells were plated assessed as above. Cell growth rates after a given treatment were normalized to the respective vehicle controls.

### Intracisternal inoculation

Animals were anesthetized with a mixture of ketamine and xylazine (100 mg/kg and 10 mg/kg, respectively) administered intraperitoneally (i/p). The fur over the head and neck was shaved and the skin was disinfected. Animals were placed in a prone position crossing a 15 ml conical tube and the head was flexed perpendicularly downwards. Ten microliters of the cell suspension (made in PBS) or of a given treatment were injected into the *cisterna magna* (between the occiput and the first cervical vertebra) using a 0.5 ml syringe equipped with a 30-gauge needle. For cancer cell inoculation, 2 × 10^3^ of LLC-LeptoM or 5 × 10^2^ of 4T1-LeptoM cells were delivered, respectively. Tumor growth was assessed by non-invasive bioluminescence imaging. For survival studies, experimental endpoint was defined as the onset of tumor-related symptoms, including, including neurological symptoms (such as seizure and paralysis), lethargy, weight loss, hunching, and macrocephaly.

### Subcutaneous and intra-lung inoculation

Mice were anesthetized with isoflurane (3-5%; Covetrus #11695067772) and the fur covering the left flank region, or the chest was shaved for subcutaneous and intra-lung cell injection, respectively. The skin was disinfected and 50 μl contained 5 × 10^4^ of control or Cxcl1 knockout LLC-LeptoM cells were injected. For subcutaneous tumors, dimensions were measured using a digital caliper (VWR) starting one week after injection of cancer cells and continued every 3 - 4 day until the tumor size reached 1 cm at its largest dimension. Tumor volume was calculated using the following formula: Tumor volume = (length × width)^2^/2. Intra-lung tumor growth was assessed by bioluminescence imaging for two weeks after injection of cancer cells.

### Irradiation procedure

Animals were anesthetized with isoflurane in a flow of oxygen (0.5 L/min). Isoflurane concentration was 3-5% for induction and 1.5% for maintenance during the irradiation procedure. Animals were placed in a prone position and irradiated using X-RAD320 irradiator (Precision; North Branford, CT, USA) applying the following setting: 250 kV; 12 mA; using 0.25 mm copper filter; distance for radiation source to the animal body: 50 cm; irradiation field: 20 × 20 cm; dose rate: 117.5 cGy/min. An adjustable lead shielding was costumed to enable irradiation of the head only (whole brain irradiation; WBI); simultaneous irradiation of the head and the spine column (craniospinal irradiation; CSI); or irradiation of the spine only (spine irradiation; SI). Depending on the experiment, animals received a total radiation dose of 10 or 20 Gy CSI, WBI or SI given hypofractionated at a dose of 2 or 4 Gy per fraction, respectively, delivered over five consecutive days. In one experiment during the optimization phase of our CSI protocol, a 10 Gy CSI was given as a single fraction. Sham controls (SH) were anesthetized for the similar duration of time required to deliver a particular radiation dose, but received 0 Gy radiation. After completion of the irradiation procedure, animals were allowed to recover from the anesthesia and returned to their cages.

To evaluate the secretion of Cxcl1 by cancer cells in response to irradiation, LLC-LeptoM were seeded in 6-well plates at a density of 5 × 10^4^ and grown on complete culture media overnight. Cells were carefully rinsed three times with PBS, and then grown in serum-free culture media. Cells were irradiated with a single dose of 4 Gy using the XRAD-320 irradiator. SH control cells were transported together with the irradiated cells and placed in the irradiation chamber for a similar duration of time required to deliver 4 Gy, but received 0 Gy radiation.

### Treatment with the Cxcr2 antagonist

Interruption of Cxcr2 signaling was perform by treating the animals with the Cxcr2 antagonist SB262510 (Tocris #2724). The antagonist was dissolved in dimethyl sulfoxide (DMSO; Corning #25950CQC), aliquots were made and stored at −20°C. To prepare the injection solution, the antagonist was further diluted in 1× PBS to bring the DMSO concentration to 10% and the antagonist concentration to 133 μM. For the i/p treatment, 200 μl of vehicle (10% DMSO) or the antagonist solution were injected per mouse, daily for 3 weeks, starting immediately after inoculation of the cancer cells. For the intracisternal treatment, vehicle or the antagonist solution were co-injected with the cancer cells and the treatment was maintained by intracisternal injection of 10 μl of vehicle or the antagonist solution starting 24 h after inoculation of cancer cells and continued every 48 h during the first week after inoculation of cancer cells, and every 72 h during the second week. For the late treatment with the SB265610 combined with craniospinal irradiation (CSI; total 10 Gy delivered as 2 Gy × 5 fractions), treatments were started one week after inoculation of cancer cells. Animals were divided into four treatment groups: Sham irradiated injected with vehicle (SH + Veh), CSI treated injected with vehicle (CSI + Veh), sham irradiated injected with SB265610 (SH + SB), or CSI treated with injected with SB265610 (CSI + SB). Treatment with vehicle or the antagonist solutions were delivered by intracisternal injection of 10 μl of either solution given every 48 for a period covering the second and third week after inoculation of cancer cells.

### Bioluminescence imaging

The cancer cell growth (tumor burden) after intracisternal or intra-lung inoculation of cancer cells was monitored using bioluminescence imaging. Animals were anesthetized with isoflurane (3-5% for induction and 1.5% for maintenance) in a flow of oxygen. Anesthetized animals were injected retro-orbitally with a solution of with D-Luciferin potassium salt (Goldbio; #LUCK; 15 mg/ml) and imaged using IVIS Spectrum-CT (Caliper Life Sciences). Images were analyzed using Living Image software (version 4.3.1). Data presented as a percentage of the increased luminescence signal as compared the day of inoculation of cancer cells (day 0) or to the signal recorded the time of starting a given treatment.

### Sampling and tissue collection

Animals were deeply anesthetized with a mixture of ketamine and xylazine. The CSF was collected from the *cisterna magna* using a 30-guage beveled needle and transferred to 1.5 ml microtube and kept on ice. The CSF was centrifuged at 600 × g for 5 min at 4°C. The cell-free supernatant was transferred into a new microtube and stored at −80°C until being analyzed using Olink target 96 mouse exploratory panel (www.olink.com). For tissue and floating cells collection, animals were transcardially perfused with 1× PBS. Brains were collected and placed on a petri dish. The brain surface, ventricles and the meninges that remained in the cranium were washed with 1× PBS to collect the non-adherent cells. The washout solution was collected into a 15 ml conical tube, placed on ice, and processed for flow cytometry. Brains were then placed into a 4% paraformaldehyde solution (FD Neurotechnologies #PF101) and stored at 4°C for 24 h, then transferred to a 30% sucrose solution (Sigma-Aldrich #S7903) made in 0.1M phosphate buffer (pH 7.4) for dehydration and processed for cryosectioning.

### Immunofluorescence staining and microscopy

The left hemisphere was sagittally cut into 50 μm free-floating sections using a sliding microtome (Thermo Scientific HM450). Serial sections of 1:9 series were generated and stored at 4°C in tubes contained a cryoprotectant solution (25% glycerol, 25% ethylene glycol in 0.1 M phosphate buffer). Sections were incubated in a citrate buffer solution (Sigma-Aldrich #C9999; pH 6.0) for 30 min at 80°C for antigen retrieval. Non-specific binding was blocked by incubating the sections in a solution of 1× Tris-buffered saline (TBS) containing 3% normal donkey serum (Jackson ImmunoResearch Laboratories #017000121) and 0.1% Triton X-100 (Sigma-Aldrich #X100) for 1 h at room temperature. Sections were then incubated with primary antibodies at 4°C for 48 - 72 h. The following primary antibodies were used; mouse anti-CXCL1 (1:500; Abcam #ab89318); goat anti-Iba1 (1:500; Abcam #ab5076); rabbit anti-Iba1 (1:1000; Wako/Fuji #01919741); chicken anti-mCherry (1:1000; Novous #NBP2-25158); rabbit anti-luciferase (1:250; Abcam #ab185924); goat anti-Icam1 (1:250; R&D Systems #AF796). After several washes with 1× TBS, sections were incubated with appropriate secondary antibodies for 2 h at room temperature. The following secondary antibodies were used; AlexaFluor-488 donkey anti-rabbit IgG (1:1000; ThermoScientific #A-21206); AlexaFluor-555 donkey anti-mouse IgG (1:1000; ThermoScientific #A-31570); AlexaFluor-594 donkey anti-chicken IgY (1:1000; Jackson ImmunoResearch Laboratories #703-585-155); AlexaFluor-647 donkey anti-mouse IgG (1:1000; ThermoScientific #A-32787); AlexaFluor-647 donkey anti-rabbit IgG (1:1000; ThermoScientific #A-31573); AlexaFluor-633 donkey anti-goat IgG (1:1000; ThermoScientific #A-21082). Hoechst 33342 (ThermoScientific #H3570) was used for nuclear counterstaining. Sections were mounted onto glass slides and coverslipped using ProLong Gold anti-fade reagent (ThermoScientific #P36930). For analysis of Cxcl1 expressing cells in leptomeninges, images were acquired using the LSM 880 Zeiss confocal scanning microscopy (Carl Zeiss, Germany) using a 40× objective lens and 1 airy pinhole setting. Z-stack images were acquired in sequential scans performed at 1 μm section intervals. Images were analyzed using Zen Blue Lite software (Carl Zeiss; version 3.3). Analysis was performed on the leptomeninges of sagittal sections leveled between 0.5 - 1.5 mm lateral bregma. Confocal representative images for Cxcl1 colocalization with phenotypic markers were acquired using 40× or 63× objective lens and 1 airy pinhole using a digital zoom of 1 or 2×.

## ELISA

The concentrations of CXCL1 in human CSF samples were assayed using the human CXCL1 ELISA kit (Abcam #ab190805). Cell-free CSF samples were thawed on ice, and assayed undiluted. Samples with concentration exceeding the highest standard provided in the kit were re-assayed diluted 1:10. The assays were performed in accordance with manufacturer’s instructions. The Cxcl1 levels in culture media were measured using the mouse Cxcl1 ELISA kit (Abcam #ab216915). The culture media were either assayed undiluted or diluted 1:100, depending on the experimental condition, and assayed in accordance with manufacturer’s instructions.

### Flow cytometry and cell sorting

Floating cells were collected from the meningeal and cranial wash were filtered through a 40 μm mesh, and centrifuged at 500 × g for 5 min at 4°C. The supernatant was discarded, and the cell pellet was suspended in 1× flow cytometry staining buffer (R&D Systems #FC001). Non-specific binding was blocked by incubating the cells with mouse fcR blocking reagent solution (Miltenyi #130-092-575). The following antibodies were used: BUV395 rat anti-mouse CD45 (Clone 30-F11; BD Horizon #564279); brilliant violet 421 anti-mouse/human CD45R/B220 (Clone RA3-6B2; BioLegend #103251); brilliant violet 650 anti-mouse NK-1.1 (Clone PK136; BioLegend #108736); brilliant violet 711 anti-mouse F4/80 (Clone BM8; BioLegend #123147); brilliant violet 785 anti-mouse Ly6G (Clone 1A8; BioLegend #127645); PerCP anti-mouse Ly6C (Clone HK1.4; BioLegend #128027); PE/Cy7 anti-mouse/human CD11b (Clone M1/70; BioLegend #101216); Alexa Fluor 647 anti-mouse CD11c (Clone N418; BioLegend #117314); APC/fire 750 anti-mouse TCR β Chain (Clone H57-597; BioLegend #109246); PE anti-mouse Cxcr2 (R&D Systems #FAB2164P). Fixable green live/dead stain (Thermo Fisher Scientific #L34969) was used to determine cell viability. Flow cytometry analysis was run on BD Fortessa (BD Biosciences). Data analysis was performed using FlowJo (Becton Dickinson). For analysis of Cxcr2 expression and sorting of LLC-LeptoM and 4T1-LeptoM cells into Cxcr2-positve and Cxcr2-negative cells, cells were stained with PE anti-mouse Cxcr2 (R&D Systems #FAB2164P) and sorted at 4°C using BD FACSAriaIII (BD Biosciences) and processed for RNA extraction. For analysis of CXCR2 and EpCAM expression in the CSF of LM patients, cells were stained with Alexa Fluor 488 anti-human EpCAM (Clone 9C4; BioLegend #324210); APC anti-human CXCR2 (R&D Systems #FAB331A); BUV395 anti human CD45 (Clone HI30; BD #563792). Flow cytometry analysis was run on Cytek Aurora (Cytek Biosciences).

### Bulk RNA sequencing

Sorted cells were lysed using RLT lysis buffer (Qiagen #79216) containing 2-mercaptoethanol (1:100; Sigma-Aldrich #M6250) and stored at −80°C. Samples were thawed on ice and RNA was isolated using RNeasy Plus Micro Kit (QIAGEN, #74034). Samples were processed for bulk RNA sequencing at the Integrative Genomic Operation core facility at MSKCC. Briefly, After RiboGreen quantification and quality control by Agilent BioAnalyzer, 2 ng total RNA with RNA integrity numbers ranging from 7.5 to 10 underwent amplification using the SMART-Seq v4 Ultra Low Input RNA Kit (Clonetech catalog #63488), with 12 cycles of amplification. Subsequently, 10 ng of amplified cDNA was used to prepare libraries with the KAPA Hyper Prep Kit (Kapa Biosystems KK8504) using 8 cycles of PCR. Samples were barcoded and run on a NovaSeq 6000 in a PE100 run, using the NovaSeq 6000 S1 (14081) or SP (14142) Reagent Kit (200 cycles) (Illumina). An average of 44 million paired reads were generated per sample and the percent of mRNA bases per sample ranged from 89.5 to 91.4 and ribosomal reads averaged 0.24%. Resulting data were processed in RStudio (version 2023.06.1-524) with DESeq2 for principal component analysis, differential gene expression analysis, and Gene Ontology analysis.

### Statistical analysis

Olink protein expression data was acquired from thirteen LM patients with multiple CSF collections after pCSI (baseline, 1-, 3-, and 6-months). For four patients, the average value of the other 9 patients for each protein was used as a baseline value. At each time point, an average of the protein expression values was calculated and the log2 fold-change was considered by dividing each protein average expression by its average baseline value. All analyses were performed using R (version 4.2.1). Other statistical analyses were performed using GraphPad Prism software (version 9.0.). Quantification data are presented as mean ± SEM. *t*-test was applied when comparing two groups and One-way ANOVA was applied when comparing more than two groups. Two-way ANOVA was used when comparing the effect of a given treatment over different time points. Tukey’s or Bonferroni’s tests were used for multiple comparisons and corrections. Log-rank (Mantel-Cox) test was used to compare survival in the mouse studies and for correlation of CXCL1 levels with overall survival and the central nervous system progression-free survival in patients. Specifications of statistical tests, the numbers of animals used, or the number of independent experiments are reported in each figure legend. Statistical significance was considered when *p* values were < 0.05.

---

**Fig. S1:**
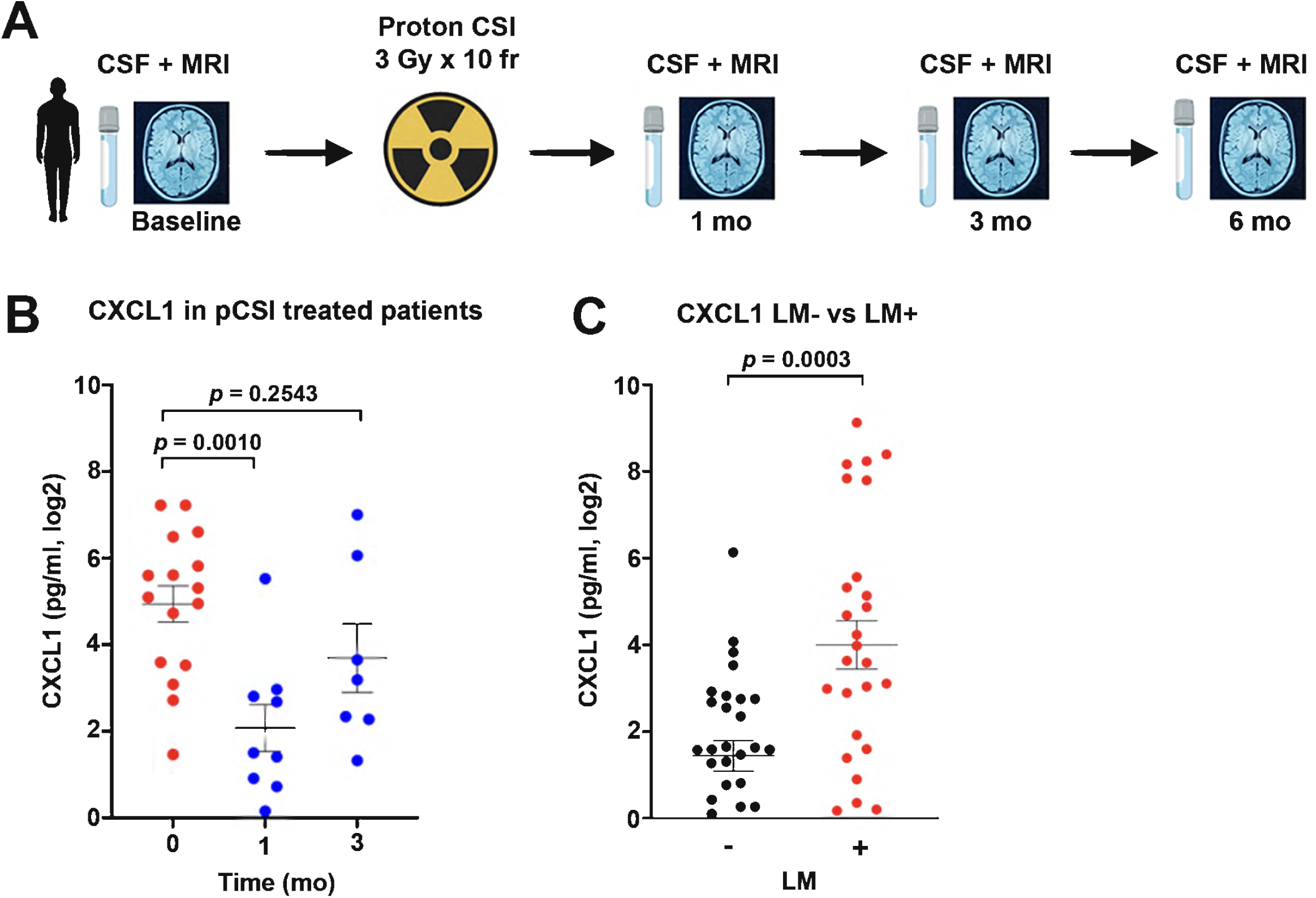
Leptomeningeal metastasis (LM) increases CXCL1 in the cerebrospinal fluid (CSF). (A) Study overview. CSF collection and magnetic resonance imaging (MRI) were performed before treatment with proton craniospinal irradiation (pCSI) (baseline; timepoint 0) and at 1-, 3- and 6-months (mo) post pCSI. Gy = Gray. Fr = fraction. (B) Dot plot illustrating ELISA measurements of CXCL1 levels in the CSF from LM patients before and after pCSI. Data represent mean ± SEM. One-way ANOVA with Bonferroni correction. Baseline (0): n = 16; 1 month: n = 9; 3 months: n = 7. Baseline vs 1-month *p* 0.0010; Baseline vs 3-month *p* = 0.2543. (C) Dot plot illustrating ELISA measurements of CXCL1 levels in the CSF from patients with and without LM, LM+ and LM-, respectively. Data represent mean ± SEM. Unpaired *t*-test. LM-n = 31; LM+ n = 27. *p* = 0.0003.

**Fig. S2:**
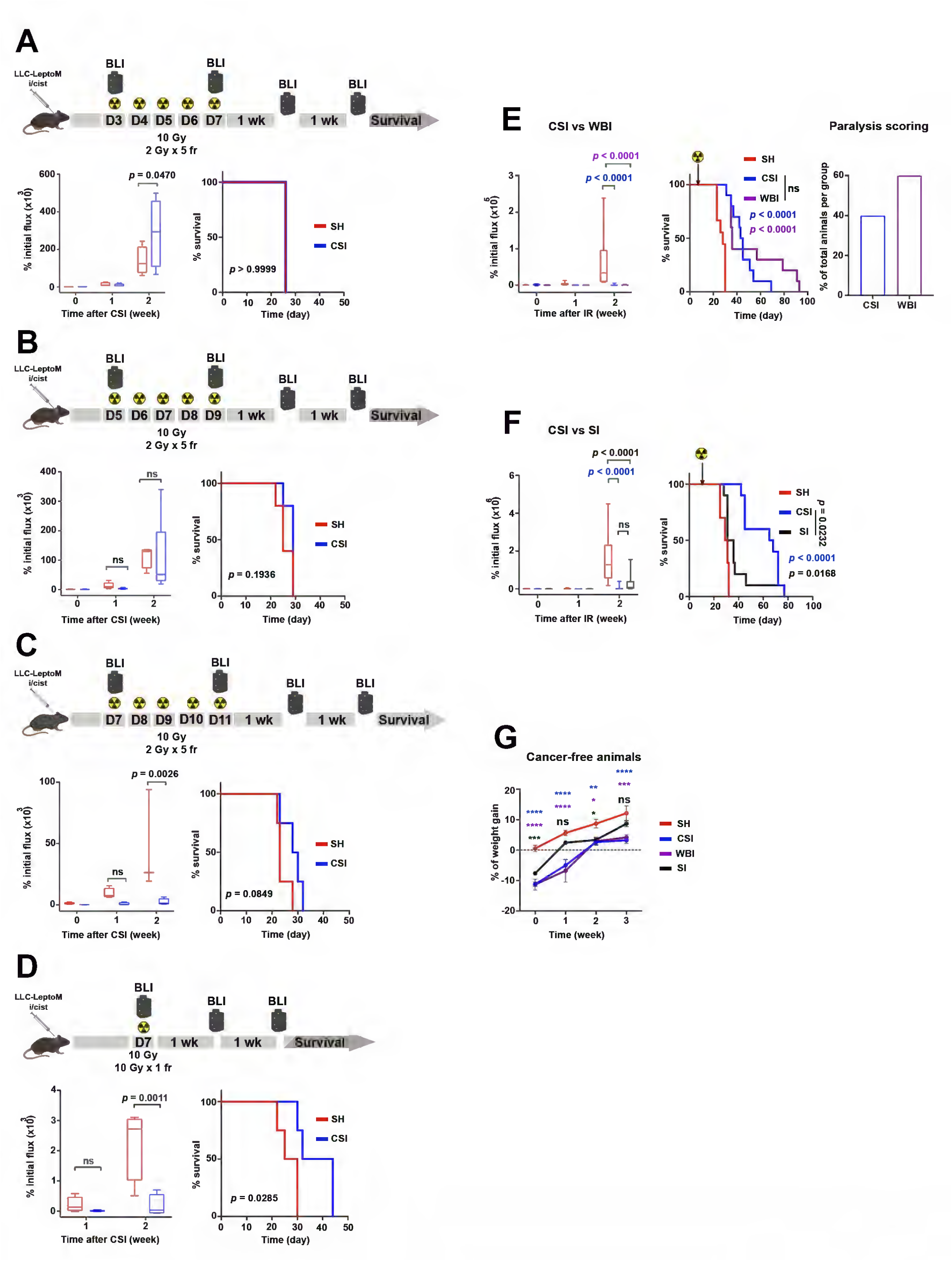
Generation of a mouse CSI model. (A) Upper panel: Scheme showing the experimental design. CSI started on day (D) 3 after intracisternal (i/cist) inoculation of LLC-LeptoM cells. BLI = Bioluminescence imaging. Wk = week. Lower left: Tumor growth quantified by BLI. Data represent mean ± SEM. Two-way ANOVA with Bonferroni correction. Sham controls (SH) n = 4; CSI n = 5. *p* = 0.0470. Lower right: Kaplan-Meier survival curve *p* > 0.9999. (B) Upper panel: Scheme showing the experimental design. CSI started on D5 after i/cist inoculation of LLC-LeptoM cells. Lower left: Tumor growth quantified by BLI. Data represent mean ± SEM. Two-way ANOVA with Bonferroni’s multiple comparisons test. SH n = 5; CSI n = 5. ns = not significant. Lower right: Kaplan-Meier survival curve. *p* = 0.1936. (C) Upper panel: Scheme showing the experimental design. CSI started on D7 after i/cist inoculation of LLC-LeptoM cells. Lower left: Tumor growth quantified by BLI. Data represent mean ± SEM analyzed using two-way ANOVA with Bonferroni’s multiple comparisons test. SH n = 4; CSI n = 4. *p* = 0.0026; ns = not significant. Lower right: Kaplan-Meier survival curve. *p* = 0.0849. (D) Upper panel: Scheme showing the experimental design. CSI with a single dose of 10 Gy delivered on D7 after i/cist inoculation of LLC-LeptoM cells. Lower left: Tumor growth quantified by BLI. Data represent mean ± SEM. Two-way ANOVA with Bonferroni correction. SH n = 4; CSI n = 4. *p* = 0.0011; ns = not significant. Lower right: Kaplan-Meier survival curve. *p* = 0.0285 (E) CSI vs whole brain irradiation (WBI). Left: Tumor growth quantified by BLI. Data represent mean ± SEM. Two-way ANOVA with Bonferroni correction. SH n = 9; CSI n = 10; WBI n = 10. SH vs CSI *p* < 0.0001; SH vs WBI *p* < 0.0001. Middle: Kaplan-Meier survival curve. SH vs CSI *p* < 0.0001; SH vs WBI *p* < 0.0001. ns = not significant. Right: Bar plot showing the percentage of animals with observed paralysis after each treatment. (F) CSI vs spine irradiation (SI). Left: Tumor growth quantified by BLI. Data represent mean ± SEM. Two-way ANOVA with Bonferroni correction. SH n = 10; CSI n = 10; SI n = 10. SH vs CSI *p* < 0.0001; SH vs SI *p* < 0.0001. ns = not significant. Right: Kaplan-Meier survival curve. SH vs CSI *p* < 0.0001; SH vs SI *p* = 0.0168; CSI vs SI *p* = 0.0232. (G) Radiation toxicity tested in cancer-free animals. Left: Weight change over time. Weight was recorded for 3 weeks after completion of irradiation. Data are represented as a percent change and represent mean ± SEM. Two-way ANOVA with Bonferroni correction. SH n = 9; CSI n = 10; WBI n = 9; SI n = 10. \**p* < 0.03; \*\**p* = 0.0088; \*\*\**p* < 0.0001; \*\*\*\**p* < 0.0001; ns = not significant.

**Fig. S3:**
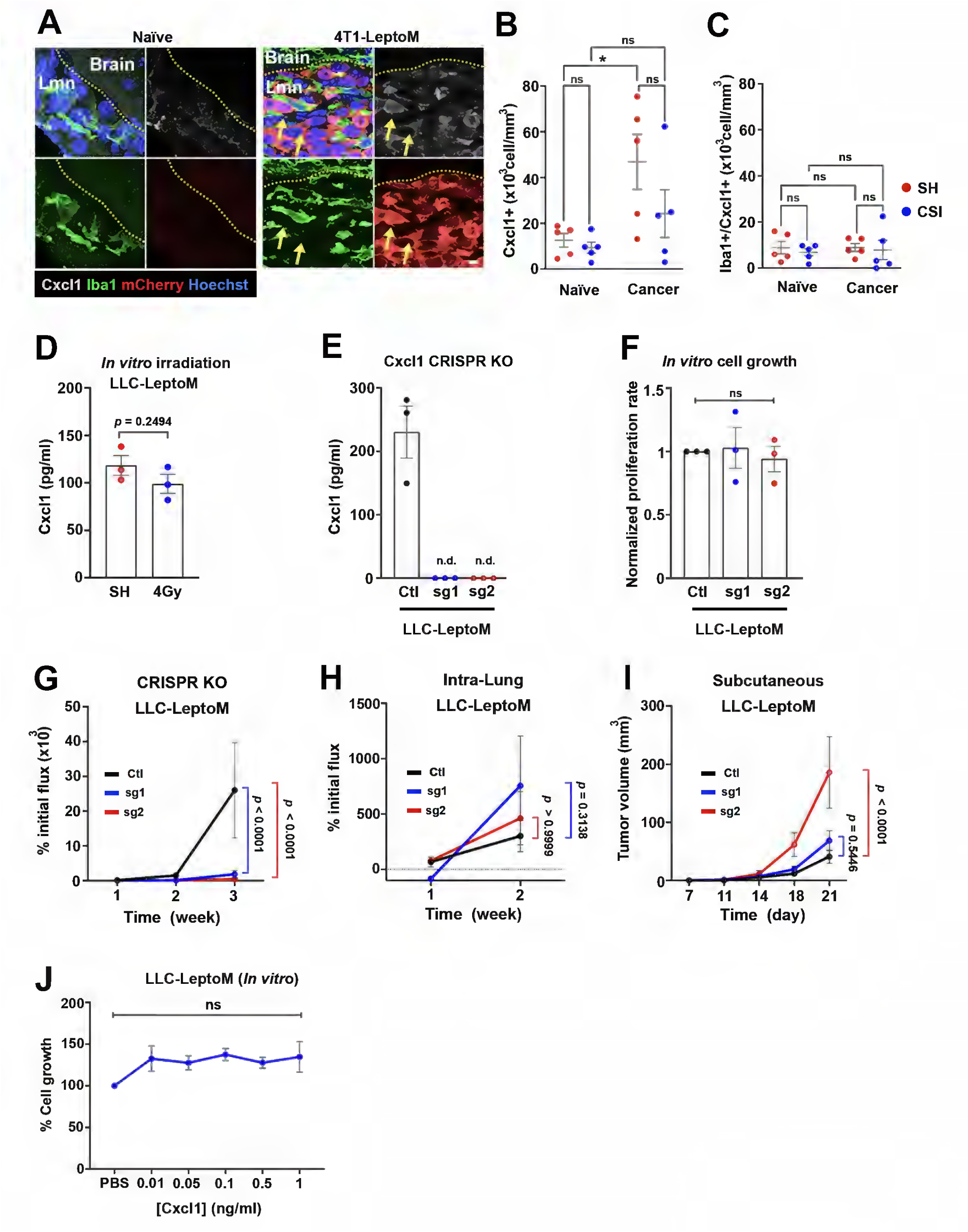
Cxcl1 expression within the LM microenvironment and its role on LM growth. (A) Confocal images showing Cxcl1 expression in leptomeninges (Lmn) of naïve and 4T1-LeptoM inoculated mice 2 weeks after injection of cancer cells. White: Cxcl1; Green: Iba1; Red: m-Cherry (expressed by cancer cells); Blue: Hoechst, nuclear counter stain. Cxcl1 expression seen in cancer cells (yellow arrows). Scale bar = 10 μm. (B) Dot plot illustrating quantitation of Cxcl1 expressing cells (Cxcl1+) in leptomeninges of naïve and 4T1-LeptoM inoculated mice 1 week after CSI. Data represent mean ± SEM. Two-way ANOVA with Tukey’s multiple comparisons test. n = 5 per group. \**p* = 0.05; ns = not significant. (C) Dot plot illustrating quantitation of Iba1+ cells co-expressing Cxcl1 (Iba1+/Cxcl1+) in leptomeninges of naïve and 4T1-LeptoM inoculated mice at 1 week after CSI. Data represent mean ± SEM. Two-way ANOVA with Tukey’s multiple comparisons test. n = 5 per group. Ns = not significant. (D) Histogram illustrating ELISA measurements of Cxcl1 levels in culture media from sham and irradiated (4 Gy) LLC-leptoM cells. Data from 3 independent experiments. Unpaired *t*-test. *p* = 0.2494. (E) Histogram illustrating ELISA measurements of Cxcl1 levels in culture media from LLC-LeptoM after Cxcl1 knockout (KO) using CRISPR/Cas9 system. Cxcl1 was detectable in culture media from control cells (Ctl), but not in culture media from the two single guide RNAs (sg1 and sg2) targeted mouse *Cxcl1*. Data are from 3 independent experiments. (F) Bar plot showing *in vitro* cell growth of Ctl and Cxcl1 KO LLC-LeptoM cells. Data from 3 independent experiments. One-way ANOVA with Bonferroni correction. ns = not significant. (G) Line graph showing tumor growth of Ctl and Cxcl1 KO LLC-LeptoM cells when inoculated i/cist into the leptomeningeal compartment quantified by BLI. Data represent mean ± SEM. Two-way ANOVA with Bonferroni correction. Ctl n = 10; sg1 n = 10; sg2 n = 9. Ctl vs sg1 *p* < 0.0001; Ctl vs sg2 *p* < 0.0001. (H) Line graph showing tumor growth of Ctl and Cxcl1 KO LLC-LeptoM cells when inoculated intra-lung (orthotopic) quantified by BLI. Data represent mean ± SEM. Two-way ANOVA with Bonferroni correction. Ctl n = 7; sg1 n = 7; sg2 n = 10. Ctl vs sg1 *p* = 0.3138; Ctl vs sg2 *p* > 0.9999. (I) Line graph showing tumor volume of Ctl and Cxcl1 KO LLC-LeptoM cells when inoculated subcutaneously. Line graphs showing tumor volume. Data represent mean ± SEM. Two-way ANOVA with Bonferroni correction. Ctl n = 10; sg1 n = 10; sg2 n = 10. Ctl vs sg1 *p* = 0.5446; Ctl vs sg2 *p* < 0.0001. (J) *In vitro* growth assay for LLC-LeptoM after treatment with recombinant mouse Cxcl1 (rmCxcl1; 0.01 ng/ml - 1 ng/ml), or vehicle (PBS). Data from 3 independent experiments. One-way ANOVA with Bonferroni correction. ns = not significant.

**Fig. S4:**
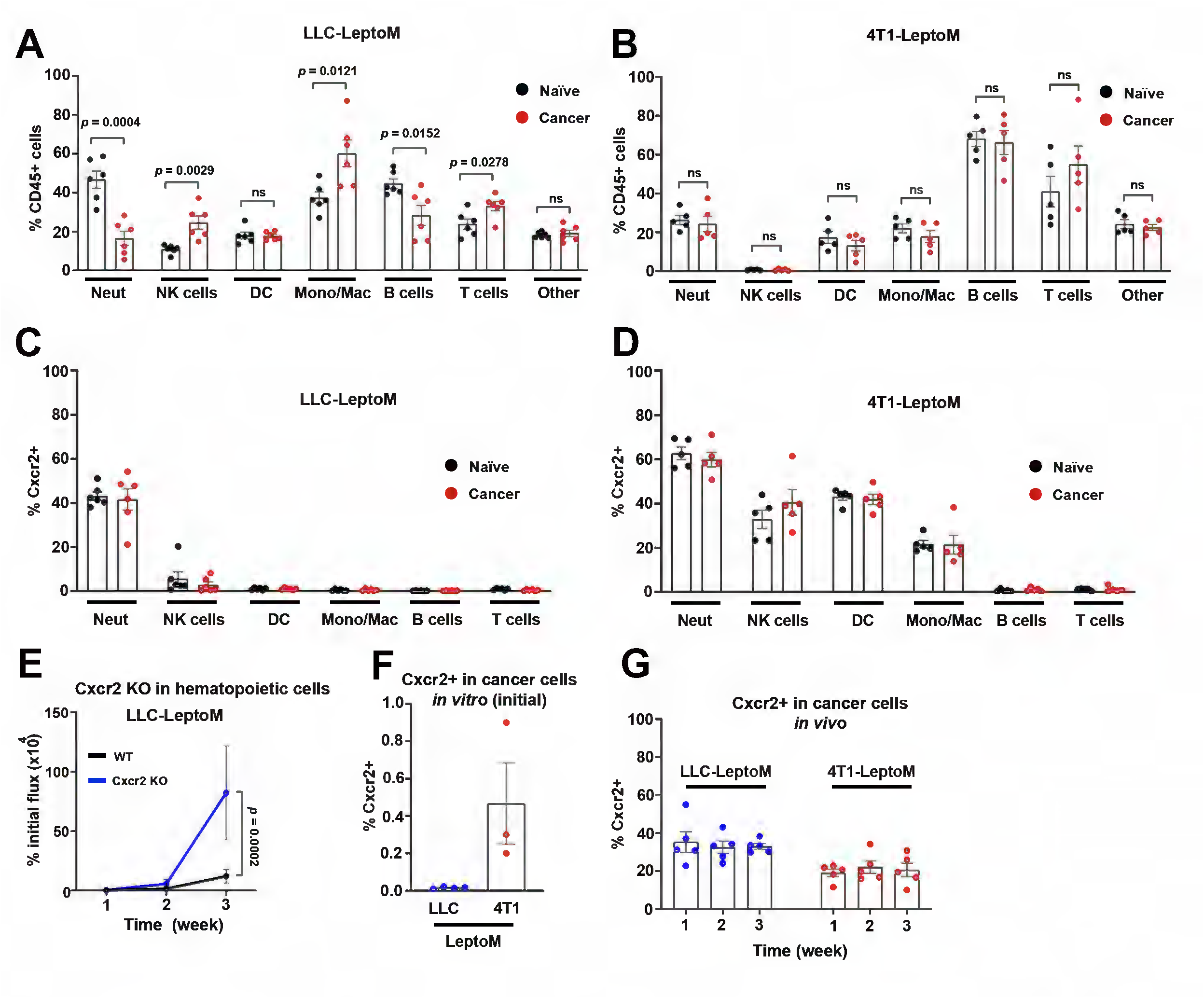
Cxcr2 expression in immune and cancer cells. (A) Histogram of immune cell types present within the LM microenvironment at 1 week after inoculation of LLC-LeptoM compared to naïve controls (received i/cist injection of PBS) quantified by flow cytometry. Data represent mean ± SEM. *t*-test. n = 6. ns = not significant. Neut = neutrophils; NK = natural killer cells; DC = dendritic cells; Mono = monocytes; Mac = macrophages. (B) Flow cytometric analysis of immune cell types present within the LM microenvironment at 1 week after inoculation of 4T1-LeptoM compared to naïve controls quantified by flow cytometry. Data are presented as percentages and represent mean ± SEM. *t*-test. n = 5. ns = not significant. (C) Histogram of Cxcr2 expressing immune cells within the LM microenvironment 1 week after inoculation of LLC-LeptoM and naive controls quantified by flow cytometry. Data represent mean ± SEM. n = 6 per each condition. (D) Immune cell Cxcr2 expression within the leptomeningeal microenvironment 1 week after inoculation of 4T1-LeptoM and naive controls as quantified by flow cytometry. Data represent mean ± SEM. n = 5 per each condition. (E) Tumor growth of wildtype (WT) and mice lacking Cxcr2 expression in immune cells (Cxcr2 KO) quantified by BLI. Data represent mean ± SEM. Two-way ANOVA with Bonferroni correction. WT n = 9; Cxcr2 KO n = 4. *p* = 0.0002. (F) Histogram of Cxcr2 expressing cells in LLC-LeptoM and 4T1-LeptoM cell lines when cells were grown *in vitro,* as quantified by flow cytometry. LLC-LeptoM: n = 4 independent experiments. 4T1-LeptoM: n = 3 independent experiments. (G) Histogram of Cxcr2 expressing cells in LLC-LeptoM and 4T1-LeptoM cell lines quantified by flow cytometry at different time points after i/cist inoculation of cancer cells. LLC-LeptoM: n = 5; 4T1-LeptoM: n = 5 per time point.

**Fig. S5:**
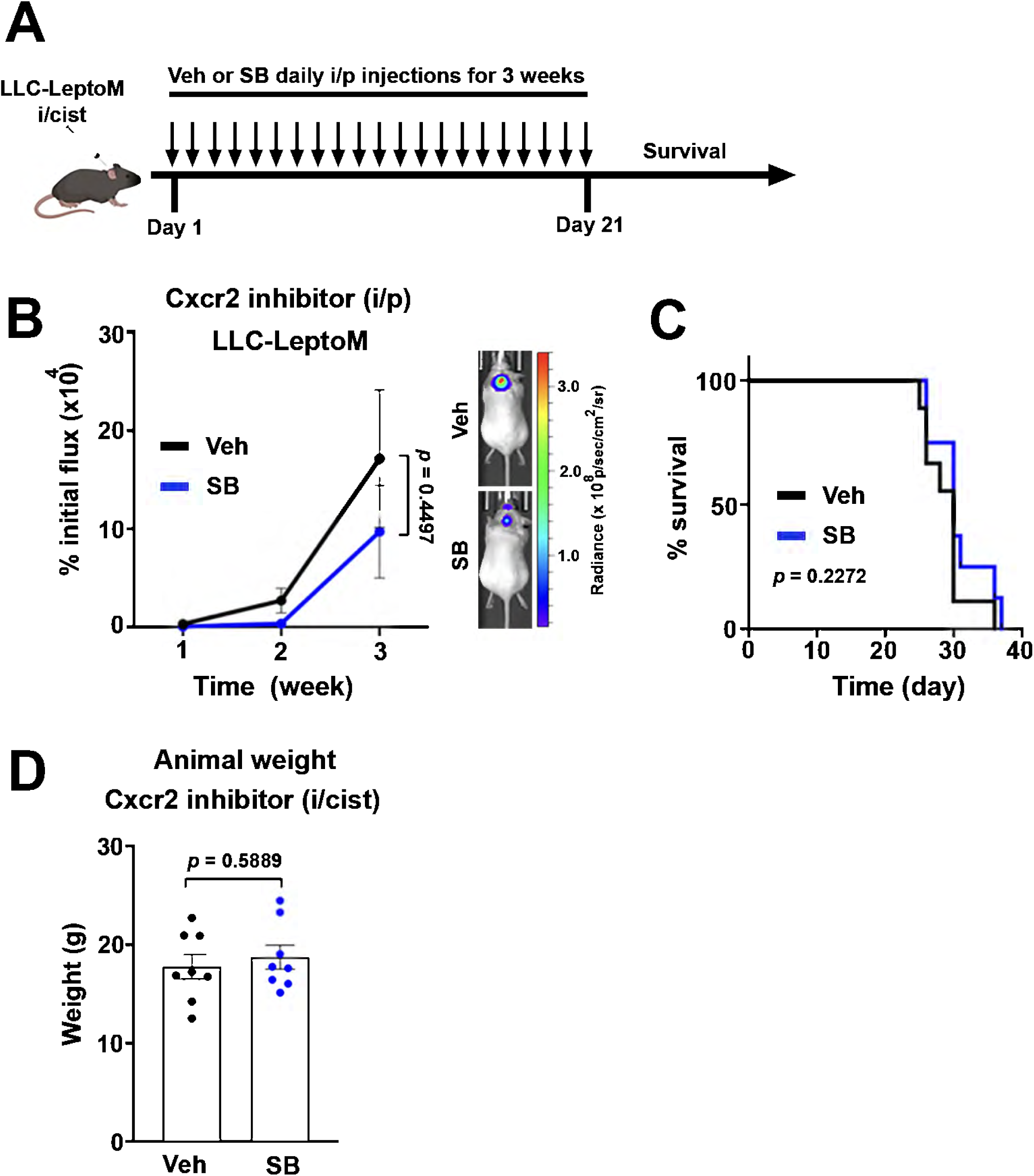
Interruption of Cxcr2 signaling. (A) Scheme for intraperitoneal (i/p) treatment with Cxcr2 inhibitor SB265610 (SB) or vehicle (Veh) in LLC-LeptoM model. Treatment started immediately after i/cist inoculation of cancer cells and maintained with daily i/p injections for 3 weeks. (B) Left: Tumor growth quantified by BLI. Right: Representative BLI images 3 weeks after inoculation of LLC-LeptoM cells. Data represent mean ± SEM. Two-way ANOVA with Bonferroni correction. Veh n = 9; SB n = 8. *p* = 0.4497. (C) Kaplan-Meier survival curve of mice in (**B**). *p* = 0.2272. (D) Animal weight recorded on the last day of treatment with i/cist injections of Veh (n = 8) or SB (n = 8). Unpaired *t*-test *p* = 0.5889.

